# Chemokines Kill Bacteria by Binding Anionic Phospholipids without Triggering Antimicrobial Resistance

**DOI:** 10.1101/2024.07.25.604863

**Authors:** Sergio M. Pontejo, Sophia Martinez, Allison Zhao, Kevin Barnes, Jaime de Anda, Haleh Alimohamadi, Ernest Y. Lee, Acacia F. Dishman, Brian F. Volkman, Gerard C.L. Wong, David N. Garboczi, Angela Ballesteros, Philip M. Murphy

## Abstract

Classically, chemokines coordinate leukocyte trafficking during immune responses; however, many chemokines have also been reported to possess direct antibacterial activity in vitro. Yet, the bacterial killing mechanism of chemokines and the biochemical properties that define which members of the chemokine superfamily are antimicrobial remain poorly understood. Here we report that the antimicrobial activity of chemokines is defined by their ability to bind phosphatidylglycerol and cardiolipin, two anionic phospholipids commonly found in the bacterial plasma membrane. We show that only chemokines able to bind these two phospholipids kill *Escherichia coli* and *Staphylococcus aureus* and that they exert rapid bacteriostatic and bactericidal effects against *E. coli* with a higher potency than the antimicrobial peptide beta-defensin 3. Furthermore, our data support that bacterial membrane cardiolipin facilitates the antimicrobial action of chemokines. Both biochemical and genetic interference with the chemokine-cardiolipin interaction impaired microbial growth arrest, bacterial killing, and membrane disruption by chemokines. Moreover, unlike conventional antibiotics, *E. coli* failed to develop resistance when placed under increasing antimicrobial chemokine pressure in vitro. Thus, we have identified cardiolipin and phosphatidylglycerol as novel binding partners for chemokines responsible for chemokine antimicrobial action. Our results provide proof of principle for developing chemokines as novel antibiotics resistant to bacterial antimicrobial resistance mechanisms.

## INTRODUCTION

The emergence of antimicrobial resistant bacteria, driven by widespread use of antibiotics and a decline in antibiotic innovation, is a major challenge to public health worldwide. Infections with antibiotic-resistant bacteria typically require prolonged hospital stays and cause higher morbidity and mortality^1,2^. A recent analysis of data from 204 countries concluded that multidrug resistant microorganisms were directly responsible for 1.27 million deaths in 2019, led by resistant strains of *Escherichia coli* (*E. coli*) and *Staphylococcus aureus* (*S. aureus*)^3^. Despite this urgent medical need, the discovery and development of new classes of antimicrobials has been relatively stagnant.

Antimicrobial peptides (AMPs) constitute a potential alternative to conventional antibiotics. AMPs are small (10-100 amino acids) and often amphipathic host-derived proteins consisting of positively charged residues that intersperse with solvent-exposed hydrophobic amino acids^4,5^. Although, like conventional antibiotics, some AMPs may disable intracellular targets, cationic AMPs are thought to kill bacteria primarily by interacting with bacterial anionic membranes^6–8^. The binding of AMPs to bacterial membranes causes membrane disorganization, increased membrane permeability, content leakage and ultimately, bacterial death^6^. Importantly, since they target non-protein structural elements fundamental for bacterial fitness and kill faster compared to conventional antibiotics, AMPs are thought to be less susceptible to bacterial resistance development^9,10^; however, the empiric evidence to support this remains limited.

AMPs constitute an important component of innate immunity and include the cathelicidin, histatin, lectin and defensin families in humans^11,12^. They promote defense against pathogenic bacteria and may also help shape the microbiome^13–15^. In addition to these professional AMPs, many members of the chemokine superfamily of chemotactic cytokines have been known for decades to have direct antimicrobial activity in vitro^16–18^. Chemokines possess many of the biochemical features of AMPs; they are small proteins (7-12 kDa), cationic, and their family-defining structure contains a C-terminal amphipathic ⍺-helix that resembles the structure of many known AMPs^19^. Of the approximately 50 different mammalian chemokines, over 20 have been shown to kill bacteria, but their antimicrobial mechanisms remain poorly understood^18,20,21^.

We recently discovered that a subset of chemokines binds with high affinity to specific anionic membrane phospholipids, including phosphatidylserine (PS) and cardiolipin (CL)^22^. PS is typically found in the inner leaflet of mammalian plasma membranes and becomes externalized during apoptosis^23–25^. CL, while present in the inner mitochondrial membrane, is absent in eukaryotic plasma membranes, but it is a common component of the plasma membrane of Gram-negative and Gram-positive prokaryotes^26,27^. We have previously demonstrated that chemokine-PS interactions may play important roles in the regulation of phagocyte recruitment for apoptotic cell clearance^22^. Here, we investigated the role of chemokine-CL interactions in bacterial killing.

## RESULTS

### The antimicrobial activity of chemokines correlates with their ability to bind phosphatidylglycerol and cardiolipin

Chemokines share many of the biochemical features of cationic AMPs and can have as potent bactericidal activity as classic AMPs^28,29^. Consistent with this, we found that human CXCL9 and CCL20 killed *E. coli* more potently than the classic AMP human beta-defensin 3 (hBD3) (Fig. 1a). This result supports that microbial killing is a bona fide activity of some chemokines, and yet the molecular properties that determine which chemokines are antimicrobial remain unknown.

**Figure 1.**
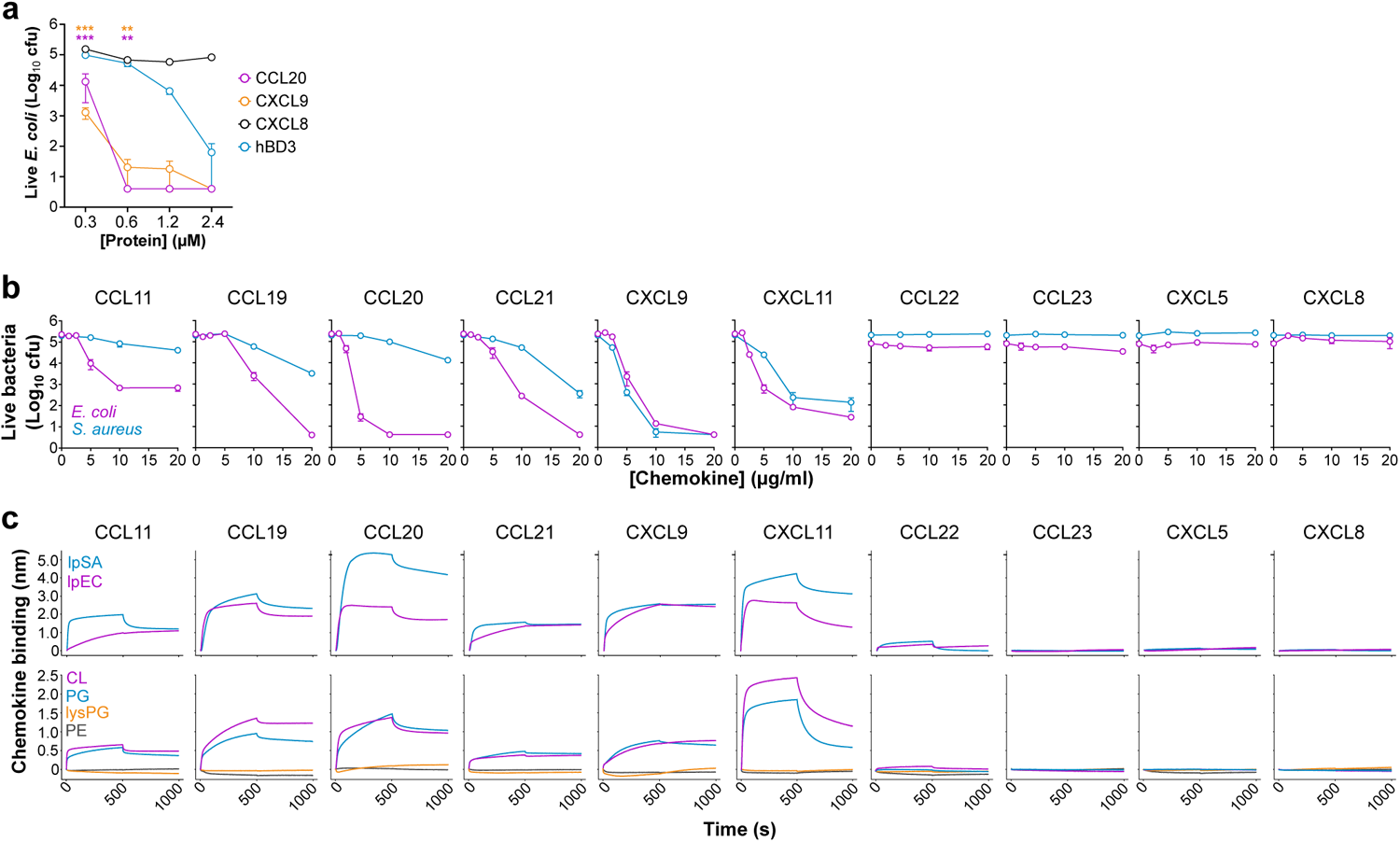
Antimicrobial chemokines bind CL and PG phospholipids. **a)** Chemokines are more potent antimicrobials than classic antimicrobial peptides. Increasing doses (x-axis) of hBD3 (blue) and the antimicrobial chemokines CCL20 (magenta) and CXCL9 (orange) were incubated with 1 x 10^+5^ cfu of *E. coli* for 2 h at 37°C. The non-antimicrobial chemokine CXCL8 (black) was used as a negative control. Samples were plated on agar plates and incubated overnight at 37°C. Data points represent the mean ± SEM cfu of 3 independent experiments analyzed in triplicate. Color-coded asterisks indicate statistically significant differences between each chemokine and hBD3 analyzed by two-way ANOVA with Dunnett’s multiple comparison test (**, p<0.01; ***, p<0.001). **b)** Antimicrobial assays. Increasing doses (x axis) of the human chemokines indicated above each graph were incubated with *E. coli* (magenta) or *S. aureus* (blue) as in panel a. Data points represent mean ± SD of the cfu counted from triplicates for each chemokine concentration. **c)** Chemokine binding to liposomes analyzed by biolayer interferometry (BLI). *Top row*, chemokine binding to liposomes replicating the phospholipid composition of *E. coli* (lpEC, magenta) or *S. aureus* (lpSA, blue). *Bottom row,* chemokine binding to phosphatidylcholine liposomes containing 30% of the phospholipids indicated on the inset of the left graph. Liposomes immobilized on BLI biosensors were incubated with 500 nM of the human chemokines indicated above each graph column. After 500 s, chemokine dissociation was recorded by incubating the biosensor with buffer alone. Lines represent the binding response in nm (y axis) over time (x axis). Data in b and c are from one experiment representative of 3 independent experiments. hBD3, human beta-defensin 3; cfu, colony formy units; CL, cardiolipin; PG, phosphatidylglycerol; lysPG, lysyl-phosphatidylglycerol; PE, phosphatidylethanolamine.

To interrogate whether the mechanism involved chemokine binding to anionic phospholipids, we first tested in a survey of 10 chemokines whether the two activities were correlated, using *E. coli* and *S. aureus* as target organisms. As shown in Figure 1b, six chemokines —CCL11, CCL19, CCL20, CCL21, CXCL9 and CXCL11— showed clear antimicrobial activity whereas four chemokines —CCL22, CCL23, CXCL5 and CXCL8—were unable to kill either organism within the tested concentration range (1.25-20 µg/ml). Of note, while CXCL9 and CXCL11 displayed similar efficacy against both organisms, CCL11, CCL19, CCL20 and CCL21 killed > 1-log colony forming units (cfu) of *E. coli* at 5-10 µg/ml but required 20 µg/ml to kill > 1-log cfu of *S. aureus* (Fig. 1b).

Next, we used biolayer interferometry (BLI) to analyze binding of the same 10 chemokines to the four main phospholipid classes found in bacterial plasma membranes: the anionic phospholipids CL and phosphatidylglycerol (PG), the cationic lysyl-PG (lysPG), and the zwitterionic phosphatidylethanolamine (PE). We tested chemokine binding to these lipids incorporated individually in liposomes of phosphatidylcholine (PC), a zwitterionic phospholipid that does not bind chemokines^22^, or combined in liposomes replicating the phospholipid composition of the plasma membrane of *E. coli* (lpEC) or *S. aureus* (lpSA), which consists of PE/PG/CL and PG/lysPG/CL, respectively, at an approximate 75/20/5 ratio in both cases^30,31^. As shown in the top row of Figure 1c, the antimicrobial chemokines CCL11, CCL19, CCL20, CCL21, CXCL9 and CXCL11 bound to both lpEC and lpSA liposomes. Furthermore, these six chemokines bound CL and PG but not PE or lysPG (Fig. 1c, bottom row). In contrast, the non-antimicrobial chemokines CCL22, CCL23, CXCL5 and CXCL8 did not bind any liposome tested (Fig. 1c). These results confirmed CL as a chemokine binding partner, identified PG as a novel lipid binding ligand for chemokines, and demonstrated a strong correlation between chemokine phospholipid-binding and antimicrobial activity, in which only chemokines that bound PG and CL were antimicrobial.

### Antimicrobial chemokines interact with *E. coli* through common bacterial binding sites and localize to anionic phospholipid-rich domains of the bacterial plasma membrane

We next tested whether bacterial membrane PG or CL could act as common binding sites for fluorophore-labeled antimicrobial chemokines. As shown in Figure 2a, the antimicrobial chemokines CXCL9, CXCL11 and CCL20, but not the non-antimicrobial CXCL8 and CCL3, were able to bind to *E. coli*^18,32^. Consistent with our hypothesis and the results in Figure 1, we found by BLI that CCL3 does not bind PG or CL (Fig. S1). Therefore, we concluded that in this survey only PG/CL-binding chemokines are able to bind the target bacteria. Furthermore, we found that the binding of a given antimicrobial chemokine to bacteria could be competed by a second antimicrobial chemokine. As shown in the representative images of Figure 2b and the quantification of the bacteria-bound CXCL11-AZ647 fluorescence intensity in Figure 2c, while the binding of CXCL11-AZ647 to *E. coli* was not affected by CXCL5 (non-antimicrobial), it was nearly neutralized when the bacteria were preincubated with the unlabeled antimicrobial chemokines CXCL11, CCL11, or CCL20. These results are consistent with a potential role of membrane CL and PG as common binding sites for antimicrobial chemokines in *E. coli*.

**Figure 2.**
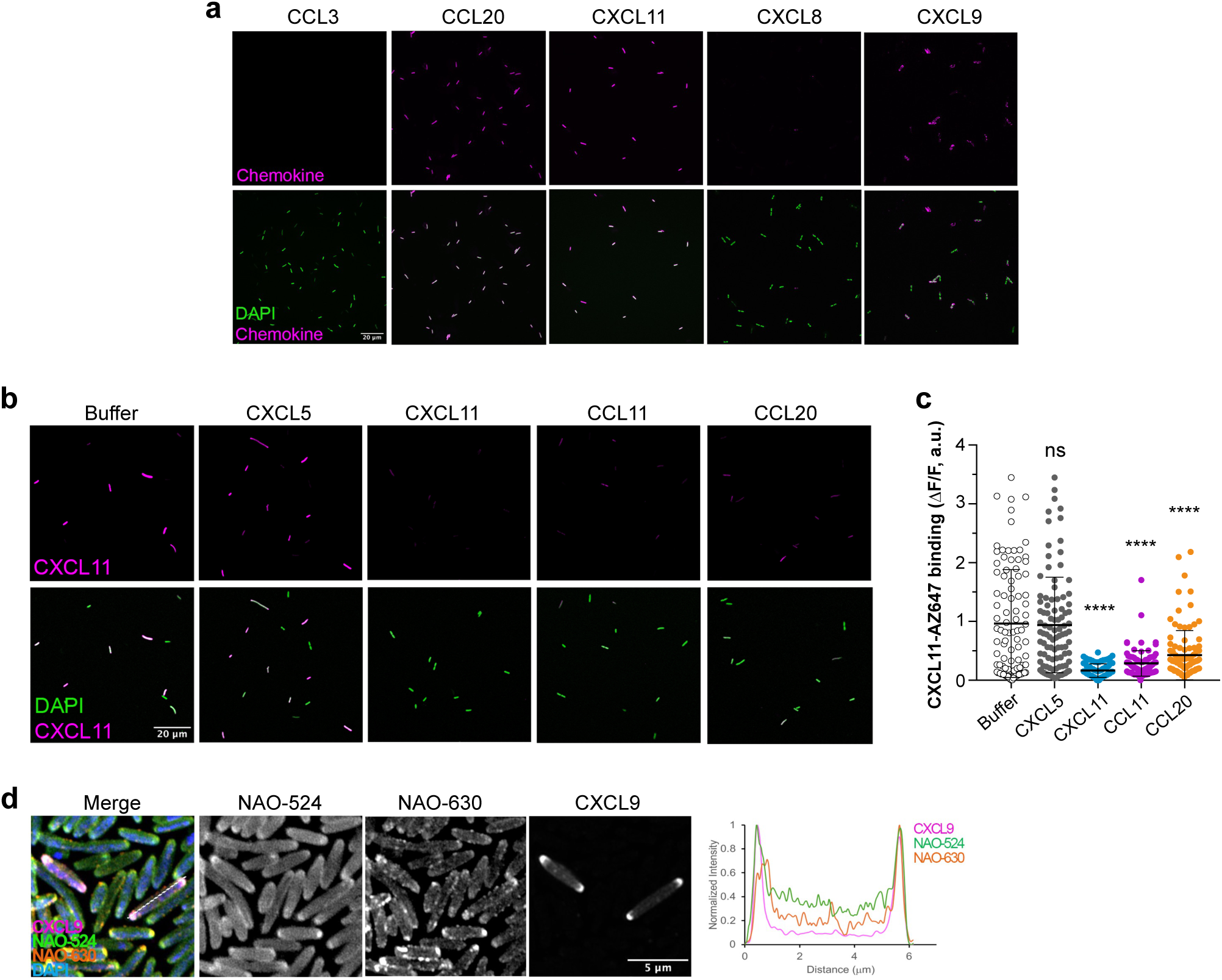
Antimicrobial chemokines share common bacterial binding sites and localize to the bacterial cell poles. **a)** Antimicrobial chemokines bind directly to bacteria. Representative images of the binding of 0.3 µM of the AZ-647-labeled antimicrobial chemokines CCL20, CCL21, CXCL9 and CXCL11 or the non-antimicrobial chemokines CCL3 and CXCL8, as indicated above each image, to *E. coli* (W3110 strain). **b and c)** Binding of antimicrobial chemokines to bacteria can be competed with other antimicrobial chemokines. In b, Representative images of CXCL11-AZ647 (0.3 µM) bound (magenta) to *E. coli* (W3110 strain) bacteria preincubated with buffer or the unlabeled chemokines indicated above each image column. In a and b, the top and bottom image rows show the staining for chemokine alone (magenta) or merged with DAPI (green), respectively. A white scale bar (20 µm) is inserted in the bottom left image. In c, quantification of the fluorescence intensity of CXCL11-AZ647 staining in bacteria preincubated with buffer alone or unlabeled chemokines as indicated on the x axis. Each dot corresponds to one bacterium (n ≈ 100). Data are from one experiment representative of 2 independent experiments. Statistical differences between each chemokine group and the “buffer” treatment group were analyzed by ANOVA with Tukey’s test for multiple comparisons (ns, not significant; ****, p < 0.0001). **d)** Antimicrobial chemokines bind to the PG/CL-rich membrane domains at the bacterial cell poles. Representative Airyscan confocal images of CXCL9-AZ647 binding to *E. coli* (W3110 strain) co-stained with NAO. Graph on the right side shows the normalized fluorescence intensity profile along a bacterium, as indicated by the dashed line in the “merge” panel, of NAO-524 (green), NAO-630 (orange) and CXCL9-AZ647 (magenta). DAPI, 4′,6-diamidino-2-phenylindole; NAO, nonyl acridine orange.

CL and PG are known to concentrate at the bacterial cell poles in the plasma membrane of *E. coli*^33^. This polar localization of CL and PG has been classically investigated by nonyl acridine orange (NAO) staining. NAO is a green fluorophore (NAO-524 [λem, max. 524 nm]) that inserts into lipid bilayers but when it binds to CL, PG or other anionic phospholipids its fluorescence emission maximum wavelength shifts to red (NAO-630, [λem, max. 630 nm])^33^. Using Airyscan confocal laser scanning microscopy and CXCL9-AZ647 as an example of a fluorescent antimicrobial chemokine, we investigated the localization of bacteria-bound chemokine in *E. coli* co-stained with NAO. As shown in Figure 2d, bacteria-bound CXCL9 concentrated and colocalized with NAO-630 at the bacterial poles. Altogether, these data indicate that antimicrobial chemokines bind to common binding sites localized at PG/CL-rich domains of the bacterial plasma membrane.

### Liposomal cardiolipin and phosphatidylglycerol protect bacteria from binding and killing by antimicrobial chemokines

To investigate the specificity of a possible CL/PG-dependent antimicrobial mechanism by chemokines, we next studied the effect of liposomes of different phospholipid compositions on the ability of chemokines to bind and kill bacteria. For this, we first tested the binding of CCL20-AZ647 to bacteria in the presence of 100 µM PC liposomes containing PE, PG or CL. As shown in the representative images in Figure 3a and the quantification of the fluorescence intensity of CCL20-AZ647 in Figure 3b, PG-and CL-containing liposomes significantly reduced or blocked, respectively, CCL20-AZ647 binding to *E. coli* whereas PE-containing liposomes had no effect. Similar results were obtained with AZ647-labelled CXCL11. Although PE reduced the binding of CXCL11-AZ647 to *E. coli* compared to the buffer treatment control, CL and PG completely blocked the binding of CXCL11 to the bacterial surface (Fig. S2). Importantly, we found that liposomes containing CL or PG, but not PE liposomes, protected *E. coli* from the antimicrobial activity of CXCL11 and CCL20 in a dose dependent manner (Fig. 3c). In particular, 100 µM of CL or PG liposomes, corresponding in these experiments to a 1:160 chemokine:lipid molar ratio, sufficed to completely inhibit the killing activity of these two antimicrobial chemokines (Fig. 3c). Furthermore, CL and PG liposomes also neutralized the killing of *S. aureus* by the antimicrobial chemokines CXCL9 and CXCL11 (Fig. 3d). These results confirmed that antimicrobial chemokines bind CL and PG and support that binding to these anionic phospholipids is required for the killing of Gram-negative and Gram-positive microorganisms by chemokines.

**Figure 3.**
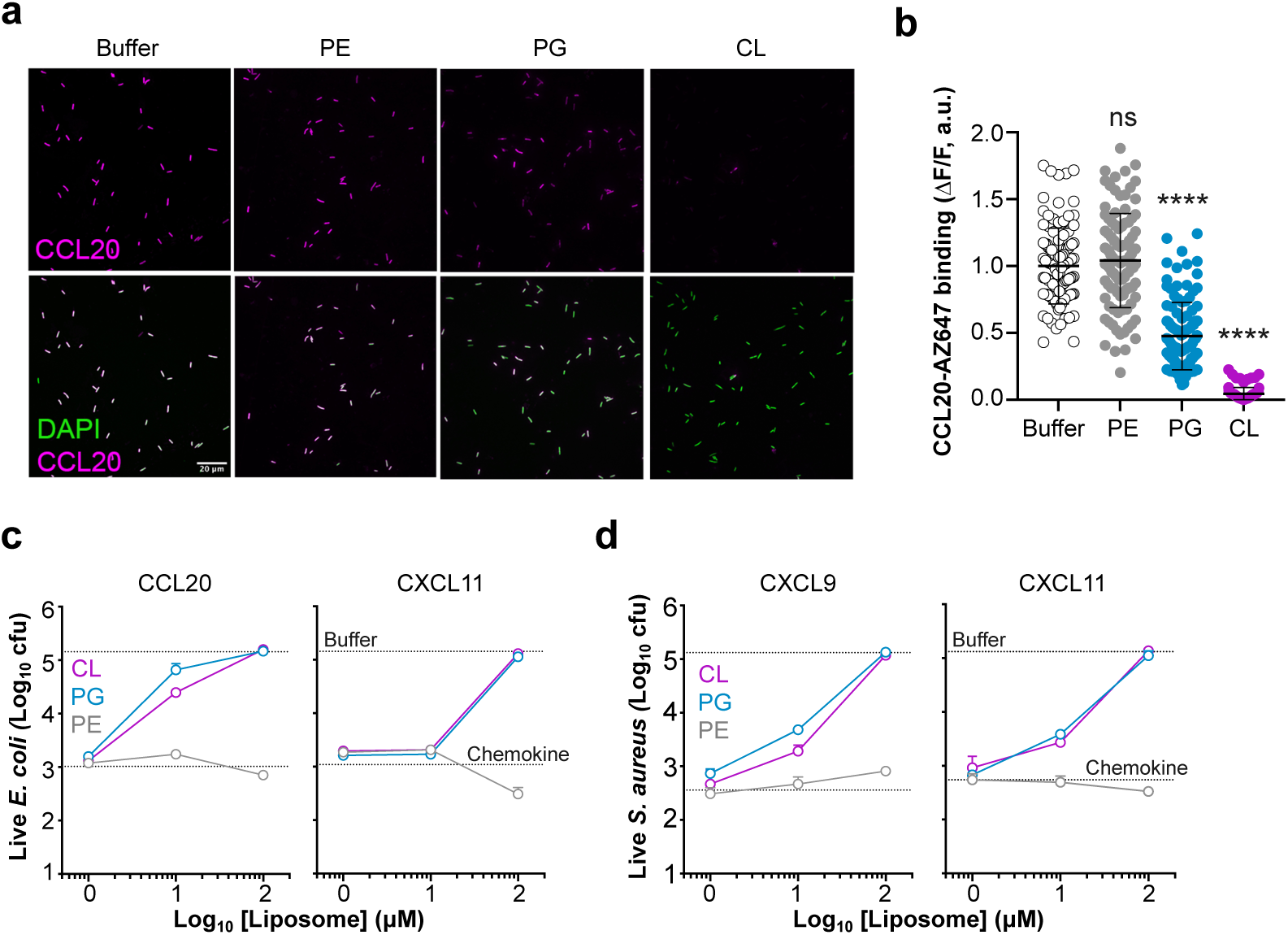
Liposomes containing CL or PG neutralize microbial binding and killing by chemokines. **a and b)** Liposomal PG and CL block chemokine binding to bacteria. In a, *E. coli* (W3110 strain) bacteria were incubated with CCL20-AZ647 in the presence of PC liposomes (100 µM) containing 30% PE, PG or CL or buffer alone as indicated above each column. Top and bottom micrograph rows show the staining for the chemokine alone or merged with DAPI, respectively. A white scale bar (20 µm) is inserted in the bottom left image. In b, quantification of the fluorescence intensity of CCL20-AZ647 bound to *E. coli* in the presence of buffer alone or liposomes containing PE, PG or CL as indicated on the x axis. Each dot corresponds to one bacterium (n ≈ 100). All liposome-treated groups were compared to “Buffer” by ANOVA with Tukey’s test for multiple comparisons (ns, not significant; ****, p < 0.0001). **c and d)** PG and CL-containing liposomes protect bacteria from antimicrobial chemokines. The chemokines (5 µg/ml) indicated above each graph were incubated for 2 h with *E. coli* (c) or *S. aureus* (d) in the presence of increasing doses (x axis) of liposomes containing 30% PE, PG or CL. Samples were plated on agar plates and cfu were counted after 18 h at 37°C. Lines represent the mean ± SD cfu from triplicates of one experiment representative of 3 independent experiments. The top and bottom horizontal dashed lines indicate the number of cfu counted when bacteria were incubated with buffer alone or with chemokine in the absence of liposome, respectively. DAPI, 4′,6-diamidino-2-phenylindole; cfu, colony forming units; CL, cardiolipin; PG, phosphatidylglycerol; PE, phosphatidylethanolamine.

### Cardiolipin-deficient bacteria are more resistant to antimicrobial chemokines

In order to further assess whether bacterial membrane phospholipids regulate chemokine antimicrobial activity, we next tested the activity of antimicrobial chemokines on the CL-deficient *E. coli* strain BKT12 compared to the wild-type parental strain W3110^34^.

Using thin layer chromatography (TLC), we first confirmed the altered phospholipid composition in the BKT12 strain. As shown in Figure 4a, BKT12 lacked CL and presented increased levels of PG compared to the parental W3110 strain. This heightened ratio of PG in BKT12 has been previously described and attributed to the deletion in this strain of the three CL synthases —ClsA, ClsB and ClsC— that consume PG to generate CL in *E. coli*^34^. However, using FACS, we found that AZ647-labeled antimicrobial CXCL9, CXCL11 and CCL21 were not only capable of binding to BKT12 bacteria but they displayed a stronger binding to BKT12 than to W3110 (Fig. 4b and 4c). Importantly, non-antimicrobial CXCL8, which does not bind CL or PG, failed to bind to either *E. coli* strain (Fig. 4b and 4c). Therefore, despite the absence of CL in BKT12, antimicrobial chemokines bind W3110 and BKT12 bacteria, which is consistent with the ability of these chemokines to bind both CL and PG phospholipids.

**Figure 4.**
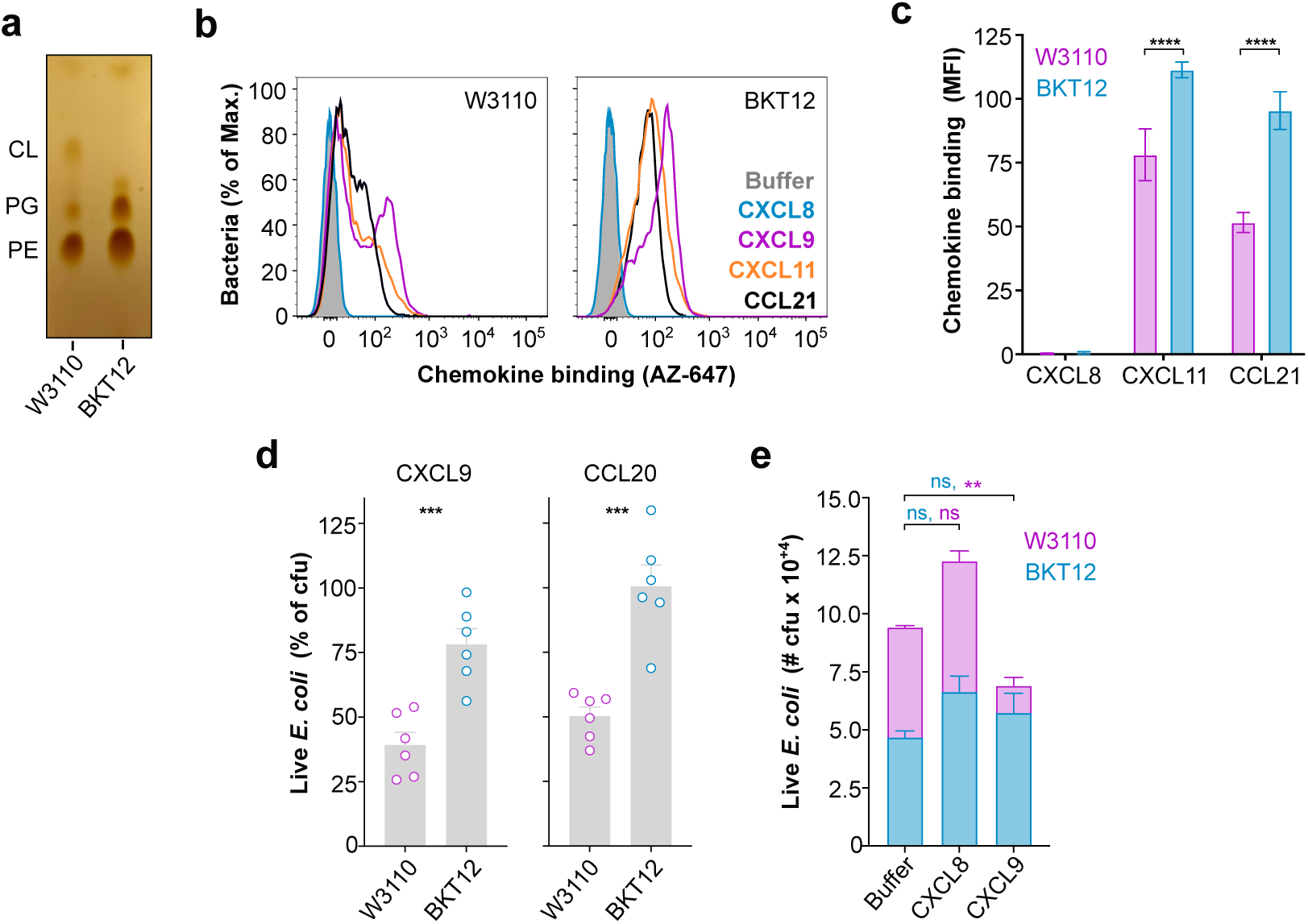
Cardiolipin-deficient bacteria are more resistant to antimicrobial chemokines. **a)** *E. coli* strain BKT12 lacks CL. TLC plate showing the phospholipid composition of *E. coli* strains W3110 and BKT12 grown to exponential phase. **b and c)** Antimicrobial chemokines bind *E. coli* W3110 and BKT12 strains. Bacteria (1 x 10^+6^ cfu) were incubated with buffer alone or AZ647-labeled chemokines (0.3 µM) and stained with SYTO24 to distinguish bacteria from debris in the flow cytometer. In b, color-coded histograms showing the binding of the chemokines indicated in the inset to W3110 and BKT12 bacteria (SYTO24^+^ events). The background signal recorded in cells incubated with buffer alone (“Buffer”) is shown with a solid gray histogram. In c, quantification of the median fluorescence intensity (MFI) of the binding of the chemokines indicated on the x-axis to W3110 and BKT12 bacteria. Data are the mean ± SD of triplicates from one experiment and are representative of 3 independent experiments. **d)** CL-deficient bacteria are more resistant to CXCL9 and CCL20. W3110 and BKT12 bacteria (1 x 10^+5^ cfu) were incubated for 2 h at 37°C with buffer or 1.2 µM of CXCL9 or CCL20 and then plated on agar plates. Surviving cfu were counted the next day and are plotted as the % relative to the number of cfu in samples incubated with buffer alone for each strain. Bars represent the mean ± SEM of data combined from 2 independent experiments. Three biological replicates were analyzed in triplicate in each experiment. Each dot corresponds to the average % cfu of one biological replicate. Data were analyzed by unpaired t test (***, p < 0.001). **e)** CXCL9 selectively kills W3110 bacteria. W3110 and BKT12 cfu were mixed 1:1 for a total of 1 x 10^+5^ cfu and incubated for 2 h at 37°C with buffer alone or 1.2 µM of CXCL8 or CXCL9. Samples were then plated in agar plates with or without kanamycin for cfu counting of total bacteria or BKT12 bacteria (kanamycin resistant), respectively. W3110 cfu were calculated as the difference between the cfu counted in agar (total bacteria) and agar-kanamycin (BKT12 bacteria) plates for each sample. Bars represent mean ± SD of 3 biological replicates analyzed in triplicate for each strain (inset) and treatment (x-axis). Data are from one experiment representative of 3 independent experiments. Differences between the buffer and the chemokine groups within each bacterial strain were analyzed by 2-way ANOVA with Bonferroni test for multiple comparisons. Statistical analysis results are color-coded and indicated above the corresponding bars (ns, not significant; **, p < 0.01). cfu, colony forming units.

To this point, we had performed all antimicrobial assays in a low-salt buffer commonly used in the field to assess AMP activity. However, we observed that CL-deficient BKT12 bacteria displayed significant levels of spontaneous death in low salt, which is consistent with the role attributed to CL in the bacterial response to osmotic stress^35^. Thus, to avoid any interfering microbial killing by osmotic shock, hereafter, all assays were performed in a 85 mM NaCl buffer, which is isosmotic to the bacterial growth media. Of note, antimicrobial chemokines are known to be sensitive to high salt^32^; however, we found that this was also applicable to the bona fide antimicrobial peptide hBD3, and that CCL20 and CXCL9 were able to kill > 50% cfu in 85 mM NaCl (Fig. S3a). Importantly, CCL20 was still as potently antimicrobial as hBD3 at this higher salt concentration, although 1.2 µM of CCL20 was required for a significant antimicrobial effect (Fig. S3b). Using these conditions, we found that 1.2 µM CXCL9 and CCL20 reduced wild type W3110 *E. coli* cfu by > 50% 2 h after treatment, whereas they killed only < 25% of the CL-deficient BKT12 strain (Fig. 4d). Furthermore, when both strains were incubated simultaneously in a 1:1 W3110:BKT12 cfu mix with chemokines or buffer alone, CXCL9 selectively killed W3110 without affecting the number of BKT12 cfu (Fig. 4E). As expected, the non-antimicrobial chemokine CXCL8 did not reduce the cfu count of either bacterial strain (Fig. 4E). We conclude that, although membrane CL is not required for the binding of antimicrobial chemokines to bacteria, chemokines are more effective antimicrobials against the parental W3110 than the CL-deficient BKT12 strain, which supports the role of bacterial membrane CL as a key molecular target for the action of antimicrobial chemokines.

### Cardiolipin mediates rapid bacteriostatic action and bacterial plasma membrane permeabilization by antimicrobial chemokines

In order to gain further insight into the chemokine antimicrobial mechanism, we next performed a FACS-based time-to-kill assay. For this, W3110 and BKT12 bacteria were treated with 1.2 µM of different chemokines and live and dead bacteria were quantified at 20, 60, 120 and 180 min after treatment by nucleic acid staining with SYTOX and SYTO24. SYTOX only permeates and stains dead bacteria with compromised plasma membrane integrity, whereas SYTO24 detects both live and dead bacteria. For reference, the antimicrobial peptide hBD3 was included in these experiments. Using this assay to quantify the number of live bacteria (SYTO24^+^ SYTOX^-^) at each time point, we found that W3110 and BKT12 treated with buffer alone grew at similar rates during the 3 h experiment, and that treatment with the non-antimicrobial chemokine CXCL8 had no effect on the growth rate of either strain (Fig. 5a). By the end of the experiment, the number of live bacteria in all samples treated with CXCL8 or buffer alone had multiplied by nearly 20-fold relative to the initial bacterial input (Fig. 5a). In contrast, all tested antimicrobial chemokines – CXCL9, CXCL11, CCL20 and CCL21 – and hBD3 stopped or significantly slowed bacterial growth (Fig. 5a). This differed from the pronounced drop in the number of live bacteria observed after incubation with 1.2 µM of the bactericidal peptide protamine (Fig. S4). Importantly, all antimicrobial chemokines stalled the growth of the parental W3110 strain to a larger extent and at earlier times than that of the CL-deficient BKT12 strain (Fig. 5a). For instance, CXCL9 and CCL20 significantly decelerated the replication of W3110 bacteria within 20 min or 1 h after treatment, respectively, whereas BKT12 bacteria treated with these chemokines grew at a rate comparable to buffer-treated BKT12 for almost 2 h (Fig. 5a). This was consistent with the different susceptibility of these two strains to CXCL9 and CCL20 after 2 h treatment shown in Figure 4d. Similarly, both CCL21 and CXCL11 inhibited W3110 replication to a greater degree than replication of the CL-deficient strain BKT12 (Fig. 5a). It is important to note that the BKT12 strain was not completely resistant to antimicrobial chemokines or hBD3. In fact, by 3 h after treatment all antimicrobial proteins were capable of effectively controlling the growth of this CL-deficient strain (Fig. 5a), possibly via their interaction with PG on the membrane of BKT12 or some other mechanism. However, these data support that membrane CL facilitates prompt control of bacterial replication by antimicrobial chemokines. Furthermore, perhaps with the exception of CXCL9 and CCL21, chemokine- and hBD3-treated bacteria seemed to overcome the initial growth retardation at later times. This highlights the importance of these time-course experiments versus the static observation obtained at one time point by cfu analysis. On the other hand, although chemokines appeared to be mainly bacteriostatic at this dose (1.2 µM), when we analyzed the presence of dead bacteria (SYTO24^+^ SYTOX^+^) at each time point, we found that W3110 was significantly more susceptible than BKT12 to direct plasma membrane permeabilization by all antimicrobial chemokines and hBD3 (Fig. 5b). As expected, dead bacteria were not found when either bacterial strain was treated with the non-antimicrobial chemokine CXCL8 (Fig. 5b). Taken together, these data indicate that chemokines can exert bacteriostatic and bactericidal effects and that both mechanisms are promoted by bacterial membrane CL.

**Figure 5.**
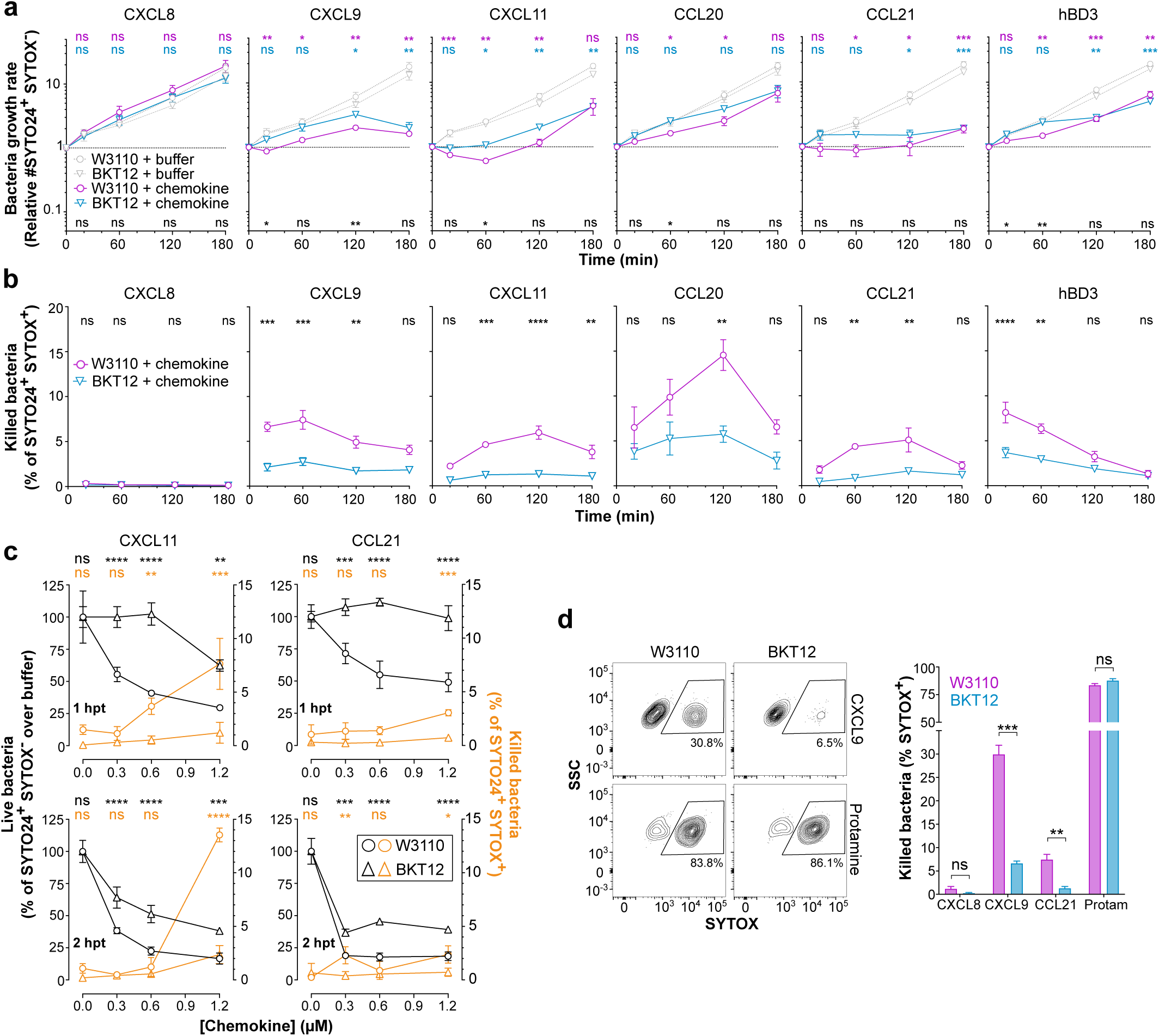
Cardiolipin promotes rapid bacteriostatic and bactericidal action by antimicrobial chemokines. **a and b)** CL is required for early bacteriostatic and bactericidal action by chemokines. FACS-based time-to-kill assays of the effect of chemokines and hBD3 on the growth (a) and killing (b) of parental W3110 and the CL-deficient *E. coli* strain BKT12. Bacteria (1 x 10^+5^ cfu) were incubated at 37°C with 1.2 µM of the protein indicated above each graph, and live (a) and dead (b) bacteria were quantified at the indicated time points (x axis) by FACS after co-staining with SYTO24 (stains live and dead bacteria) and SYTOX-Orange (stains only dead bacteria). In a, the growth ratio was represented as the ratio between the number of live bacteria (SYTO24^+^ SYTOX^-^) detected at each time point and the initial number of live bacteria recorded immediately before the start of the incubation (time 0 min) with buffer alone or the corresponding protein (as indicated in the inset of the “CXCL8” graph). In b, the % of dead bacteria (SYTO24^+^ SYTOX^+^) relative to the number of total bacteria (SYTO24^+^) for each strain (as indicated in the inset of the “CXCL8” graph) and at each time point is shown. In a and b, data are summarized as the mean ± SEM of 3 independent experiments combined with 3 biological replicates performed in each experiment. Data were analyzed by two-way ANOVA with Bonferroni test for multiple comparisons (ns, not significant; *, p < 0.05; **, p < 0.01; ***, p < 0.001). Results of statistical analyses for the comparison of buffer vs chemokine/hBD3 for each bacterial strain are color-coded and indicated above each graph. Results of statistical analyses for the comparison of W3110 vs BKT12 for each protein treatment are indicated above each x axis in black in panel a and above each graph in panel b. **c)** CL facilitates rapid bacteriostatic effects by low doses of antimicrobial chemokines. W3110 and BKT12 bacteria (as indicated in the inset of the bottom right graph) were incubated with decreasing doses (x axis) of CXCL11 and CCL21 and the number of live and dead bacteria were analyzed 1 or 2 h post treatment (hpt) (as indicated on the left of each graph row) by FACS, as in a and b. The number of live bacteria (left y-axis, black lines) detected at different chemokine concentrations is represented as % relative to the number of live bacteria recorded in the buffer-treated group for each strain and time point. The % of dead bacteria (right y-axis, blue lines) for each chemokine concentration and bacterial strain was calculated as in panel b. Data are shown as mean ± SD of three biological replicates from one experiment representative of 3 independent experiments. Data were analyzed by 2-way ANOVA with Bonferroni test for multiple comparisons. Results of the statistical analysis of the bacteriostatic (black lines) and killing activity (blue lines) for the W3110 vs BKT12 comparison are color-coded and indicated above each graph (ns, not significant; *, p < 0.05; **, p < 0.01; ***, p < 0.001; ****, p < 0.0001). **d)** CL is required for direct killing by antimicrobial chemokines. W3110 and CL-deficient BKT12 bacteria were incubated with buffer alone or 4.8 µM of CXCL9, CCL21 or protamine (Protam), and dead bacteria (SYTOX^+^) were analyzed 20 min after treatment by FACS, as in panel b. On top, representative contour plots showing the % of SYTOX^+^ cells detected after treatment of W3110 or BKT12 bacteria (as indicated above each plot column) with CXCL9 or protamine (as indicated on the right of each plot row). At the bottom, bar graph showing the quantification of the % of SYTOX+ W3110 or BKT12 bacteria (as indicated in the inset) in each treatment group. Bars show the mean ± SD of triplicates from one experiment representative of 3 independent experiments. Data were analyzed by two-way ANOVA with Bonferroni test for multiple comparisons (ns, not significant; **, p < 0.01; ***, p < 0.001). hBD3, human beta-defensin 3; hpt, hours post treatment; Protam, protamine.

One caveat to our interpretation of the time-to-kill data is that in these experiments the bacteriostatic and bactericidal activities may influence each other. For instance, the bactericidal effect may reduce the total number of live bacteria and as a result decelerate the growth of the overall population. Although the fact that the most potent bacteriostatic chemokine (CCL21) was not also the most potent bactericidal chemokine (CCL20 was) suggested that this bacteriostatic-bactericidal interrelation might not have a significant impact in our experiments, we next attempted to uncouple these two mechanisms to clearly resolve the role of membrane CL in the antimicrobial activity of chemokines. For this, we first performed time-to-kill assays with decreasing amounts of chemokine aiming to detect bacteriostatic effects at low non-bactericidal doses of chemokine. As shown in Figure 5c, and consistent with the data in Figure 5b, high doses of CXCL11 and CCL21 (> 0.6 µM) displayed detectable and significant levels (up to 14% SYTOX^+^ cells) of direct killing activity (orange lines) against W3110 but only marginal bactericidal activity (< 2% SYTOX^+^ cells) against the CL-deficient BKT12 strain at 1 and 2 h post treatment (hpt). Interestingly, subthreshold concentrations for bacterial killing (0.3 µM) of CXCL11 and CCL21 reduced the number of live W3110 bacteria (gray lines) to 50% and 70%, respectively, relative to the buffer-treated group 1 hpt whereas they did not impair the growth of the BKT12 strain at this early time point (Fig. 5c, top panels). Of note, although the bacteriostatic effect against BKT12 of both chemokines at all doses became apparent 2 hpt, the growth of W3110 was still more severely reduced at this later time (Fig. 5c, bottom panels). Therefore, while not categorically required for antimicrobial chemokines to control the growth of *E. coli* eventually, we concluded that bacterial membrane CL facilitates rapid onset of chemokine-mediated bacteriostatic effects.

On the other hand, to confirm the role of CL in bacterial killing by chemokines without interference of their bacteriostatic effects, we next analyzed the number of dead W3110 and BKT12 bacteria shortly after treatment with a high dose of antimicrobial chemokines. For this, bacteria were incubated with 4.8 µM of CXCL9, CCL21, CXCL8 or the bactericidal peptide protamine, and the % of SYTOX^+^ cells were analyzed 20 min after treatment by FACS. As shown in Figure 5d, CXCL9 killed approximately 30% of W3110 bacteria but only 6% of BKT12, whereas CCL21 killed 8% and 1%, respectively. In contrast, protamine was equally effective against both strains and killed about 85% of W3110 and BKT12 bacteria (Fig. 5d), proving that the BKT12 strain was not inherently more resistant to membrane permeabilization by AMPs. As expected, CXCL8 did not kill either bacterial strain. At this early time, CXCL9 killed considerably more W3110 bacteria than CCL21; however, it is important to note that, as shown in Figure 5B, these two chemokines appear to kill with different kinetics (peak killing at ∼20 min or 2 h after treatment, respectively). Accordingly, when we analyzed the % of SYTOX^+^ bacteria 90 min after treatment, the ratio of W3110 bacteria directly killed by CCL21 increased to 14% whereas it killed only 3% of BKT12 (Fig. S5). These results demonstrate that the CL-deficient BKT12 strain is more resistant to plasma membrane permeabilization by antimicrobial chemokines.

### Antimicrobial chemokines induce a class of membrane curvature necessary for pore formation and lyse bilayer membranes containing anionic phospholipids in a cardiolipin-promoted manner

To confirm the direct membrane lytic activity of chemokines in a simpler and more direct system, we next performed liposome calcein leakage assays. In these assays, the fluorescent dye calcein is self-quenched when trapped at high concentrations inside liposomes, but it fluoresces when released and diluted into the extra-liposomal media after a membrane active peptide ruptures the liposomal membrane. Consistent with their bactericidal activity, we found that CCL19, CCL21, CXCL9 and CXCL11 as well as protamine were able to lyse liposomes that replicated the phospholipid composition of W3110 bacteria (PE/PG/CL, 75/20/5 mass ratio) (Fig. 6a). In contrast, the non-antimicrobial chemokines CXCL8 and CCL5 had no effect on the permeability of these liposomes (Fig. 6a). We have previously demonstrated that CCL5 does not bind CL or PG^22^, and consistent with our previous data, unlike CCL19, CCL5 is innocuous to bacteria (Fig. S6a).

**Figure 6.**
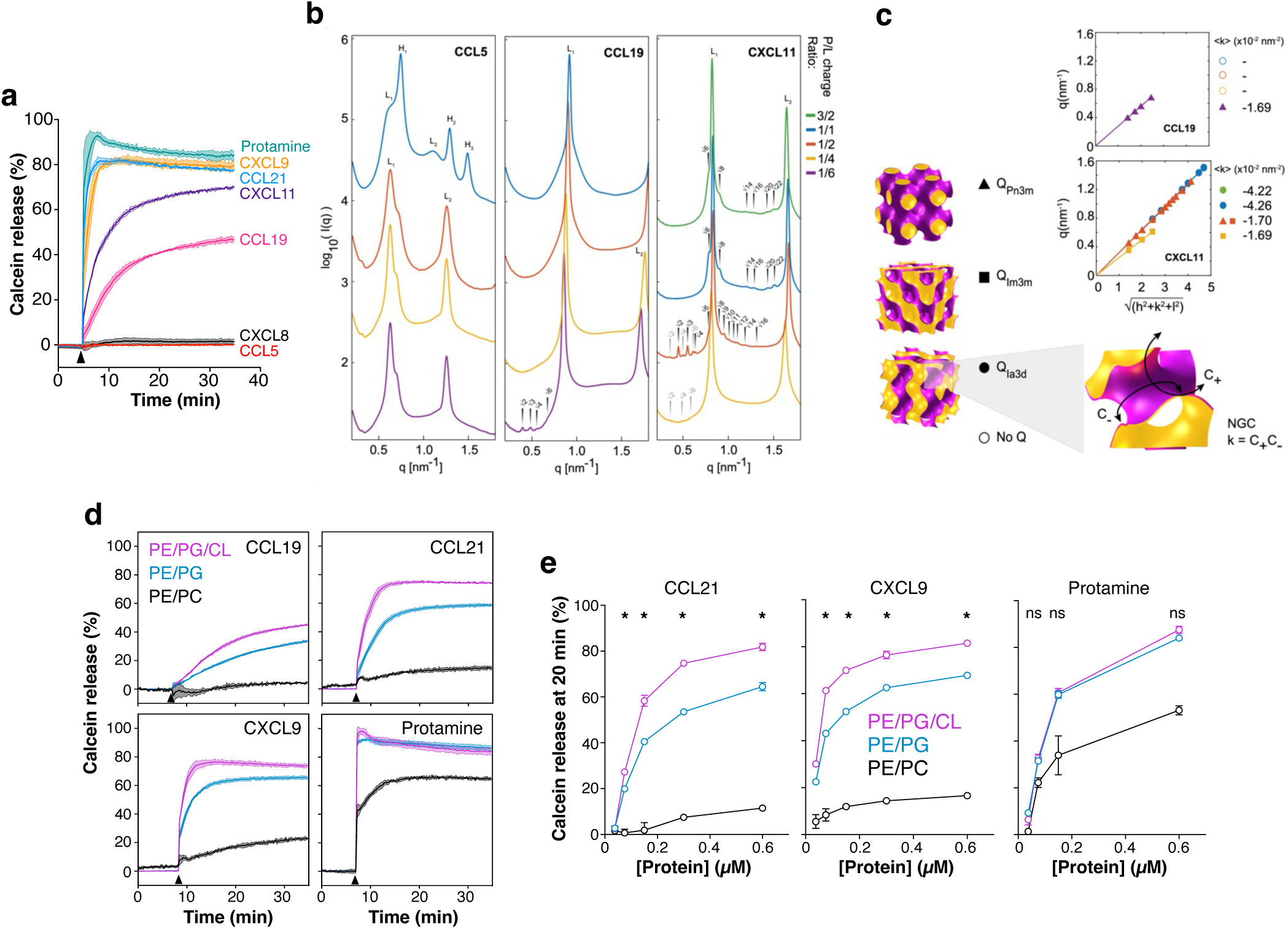
Antimicrobial chemokines lyse phospholipid bilayers in an anionic phospholipid-dependent manner. **a)** Antimicrobial chemokines disrupt liposomal membranes that mimic the phospholipid composition of the *E. coli* plasma membrane. Calcein-leakage assay showing the release of calcein from PE/PG/CL liposomes upon injection (arrowhead) of 1.2 µM of the indicated chemokines or protamine. Curves represent the % of calcein released over time (x axis) by each protein relative to the maximum calcein release observed after incubation of the liposomes with 0.1% Triton X-100. Solid lines represent the mean of 3 biological replicates. Colored shaded area around the lines represents the SD. Data are from one experiment representative of 3 independent experiments. **b)** Radially integrated SAXS spectra of chemokines CCL5, CCL19 and CXCL11 interacting with 20:80 PG:PE liposomes at neutral pH. A peptide-to-lipid (P/L) charge ratio scan from 1/6 to 3/2 was performed as indicated for each chemokine. Bragg structure peaks were indexed for the observed phases. CCL19 and CXCL11 formed cubic phases (Q) (arrowed indices). All spectra showed a lamellar phase (L), and CCL5 displayed inverted hexagonal phases (H). To facilitate visualization, spectra have been manually offset in the vertical direction by scaling each trace by a multiplicative factor. **c)** Linear fits of the peak position indexations for the cubic phases indexed in b. *Pn3m*, *Im3m* and *Ia3d* cubic phases were observed; illustration of geometric surfaces next to symbol key. Estimation of mean NGC, <K=, from linear fits is displayed next to each plot. Each color represents a P/L ratio as noted in b. **d)** Anionic phospholipids are required for the membrane lytic activity of antimicrobial chemokines. Calcein-leakage assays showing the release of calcein from 3 different types of liposomes, PE/PG/CL, PE/PG, and PEPC, as indicated in the inset of the CCL19 panel, upon injection of 1.2 µM of the proteins indicated in the inset of each graph. Data were analyzed and graphed as in a and correspond to 3 biological replicates from one experiment representative of 3 independent experiments. **e)** CL facilitates membrane disruption by antimicrobial chemokines. Calcein release from 3 different types of liposomes (as indicated in the inset of the CXCL9 graph) at 20 min after addition of increasing doses (x axis) of the proteins indicated above each graph. Calcein leakage assays were performed as in d and the % of released calcein at 20 min after protein injection was recorded. Data are represented as mean ± SD of 3 biological replicates from one experiment representative of 3 independent experiments. Data were analyzed by two-way ANOVA with Tukey’s test for multiple comparisons. Statistical differences between the PE/PG/CL and the PE/PG liposome groups are indicated above each concentration point (ns, not significant; *, p < 0.05). CL, cardiolipin; PG, phosphatidylglycerol; PE, phosphatidylethanolamine; PC, phosphatidylcholine; NGC, negative gaussian curvature.

To assess the ability of antimicrobial chemokines to disrupt membranes in a manner analogous to pore formation processes employed by membrane active cationic AMPs^36,37^, we performed high resolution synchrotron small X-ray scattering (SAXS) experiments to characterize the membrane remodeling of the chemokines on bilayer membranes. In agreement with their antimicrobial activity, we found that CCL19 and CXCL11 remodeled spherical PG-containing liposomes into negative gaussian curvature (NGC) rich cubic phases (Fig. 6b and 6c). Such membrane remodeling geometry is necessary for the restructuring of the membrane surface during pore formation, characteristic of cationic amphipathic AMPs^36,37^. Furthermore, consistent with its stronger liposomal membrane lytic activity (Fig. 6a), CXCL11 induced stronger NGC curvatures than CCL19, up to 4.26 x 10^-2^ nm^-2^ (Fig. 6c). In contrast, CCL5, a non-antimicrobial chemokine, did not induce NGC surface remodeling (Fig. 6b). Using estimates based on membrane mechanical elasticity for a typical membrane (charge density = -0.05 As/m2, Debye length = 1 nm, and line tension = 10 pN), we can infer from the SAXS measurements that the negative Gaussian curvature induced by CXCL11 and CCL19 corresponds to the formation of a transmembrane pore with a diameter of 2.4 - 2.9 nm and 2.4 - 2.5 nm, respectively^38^.

To understand the importance of each bacterial phospholipid for the membrane disrupting action of antimicrobial chemokines, we next performed liposome calcein leakage assays with PE/PG/CL liposomes (75/20/5, mass ratio), CL-lacking PE/PG liposomes (75/25, mass ratio), which mimic the phospholipid composition of the plasma membrane of the BKT12 *E. coli* strain, and PE/PC (75/25, mass ratio) liposomes lacking all anionic phospholipids. As shown in Figure 6d, 1.2 µM of CCL19, CCL21 and CXCL9 released only minimum levels of calcein from PE/PC liposomes and permeabilized PE/PG/CL liposomes more effectively than PE/PG liposomes. In contrast, protamine was equally effective at releasing calcein from PE/PG/CL and PE/PG liposomes (Fig. 6d). Similar results were obtained with a much lower dose (0.15 µM) of antimicrobial protein (Fig. S6b). We confirmed these observations in dose-response calcein leakage assays using the three different types of liposomes. Increasing doses of CXCL9 and CCL21 failed to permeabilize PE/PC liposomes and released significantly lower levels of calcein from PE/PG than from PE/PG/CL liposomes, whereas protamine leaked comparable levels of calcein from PE/PG and PE/PG/CL liposomes at all doses (Fig. 6e). Hence, the presence of CL in the liposomes was irrelevant for the membrane lytic activity of protamine whereas it facilitated membrane disruption by chemokines. These results aligned with our findings that W3110 and BKT12 are equally susceptible to membrane permeabilization by protamine, but the CL-deficient strain is more resistant to plasma membrane disruption by antimicrobial chemokines (Fig. 5d). We conclude that the membrane lytic activity of chemokines requires the presence of anionic phospholipids, particularly CL.

### Bacteria fail to develop resistance against antimicrobial chemokines

It is thought that AMPs are less susceptible to antimicrobial resistance due to their rapid action and membrane-attacking mechanism^10^. However, this has been experimentally demonstrated for very few peptides and not for chemokines. Since our data support that antimicrobial chemokines act quickly and target bacterial membranes, we next investigated whether bacteria develop resistance to antimicrobial chemokines.

For this, we purified a C-terminally His-tagged form of CCL20 (CCL20-His), which was refolded from insoluble and unfolded protein obtained by expression in bacteria (Fig. 7a). We first characterized the lipid-binding properties of CCL20-His by BLI and confirmed that this chemokine killed bacteria. As shown in Figure 7b, like untagged CCL20 (Fig. 1c), CCL20-His bound PG and CL but not PE. Then, we calculated the minimum inhibitory concentration (MIC) of CCL20-His against the wild-type *E. coli* strain W3110 by the microdilution method. Bacteria were incubated with increasing concentrations of CCL20-His in Mueller-Hinton broth (MHB), and bacterial viability was assessed the next day by luminometric determination of the ATP content in each sample. We found that the MIC of CCL20-His was 20 µM (Fig. 7c), which is in line with the MIC of other AMPs and chemokine-derived peptides^39–41^. These results confirmed that CCL20-His was an active antimicrobial chemokine.

**Figure 7.**
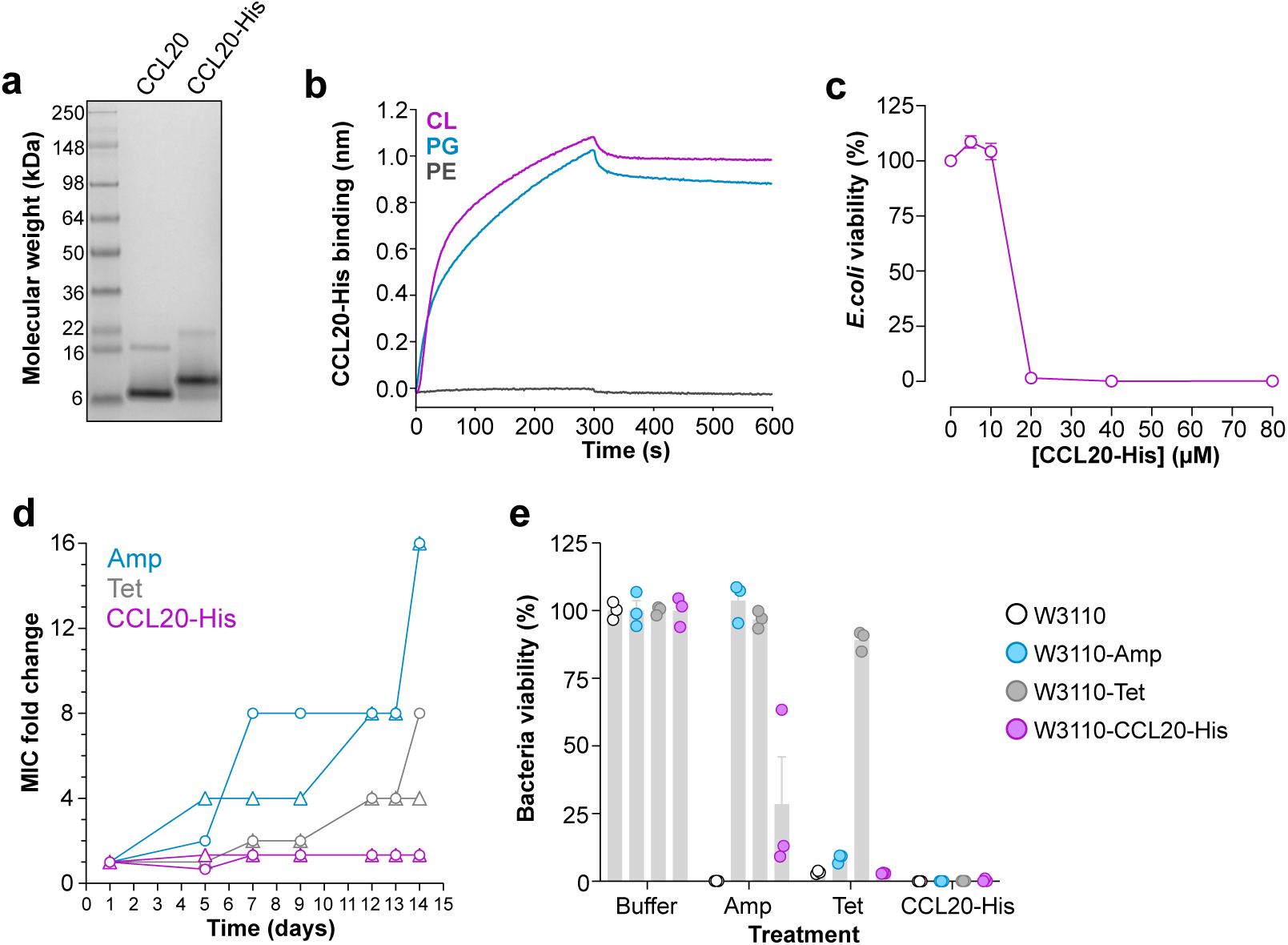
CCL20 kills wild type and antibiotic-resistant *E. coli* without triggering chemokine resistance. **a)** Coomassie Blue-stained gel (4-12% Bis-Tris) showing the SDS-PAGE analysis of commercially available CCL20 (Peprotech) and in-house produced CCL20-His. **b)** The His-tag of recombinant CCL20-His does not interfere with chemokine binding to PG and CL. BLI assays showing the binding of 0.5 µM CCL20-His to PC liposomes containing 30% of the phospholipids indicated in the inset. **c)** Microbroth dilution method for the determination of the MIC of CCL20-His. W3110 bacteria (5 x 10^+5^ cfu/ml) were incubated with increasing doses of CCL20-His in MHB and the % viable bacteria after 18 h incubation at 37°C was calculated using the BacTiter-Glo Kit. Graph shows mean ± SEM % of the bacterial viability at each chemokine concentration relative to bacteria treated with buffer alone. Data are combined from 3 independent experiments. **d)** Bacteria develop resistance against conventional antibiotics but not against CCL20-His. W3110 bacterial cultures (5 x 10^+5^ cfu/ml) were maintained in MHB for 2 weeks in the presence of a sublethal dose (0.5 x MIC) of three different antimicrobial compounds: tetracycline (Tet), ampicillin (Amp) or CCL20-His, as indicated on the inset. On selected dates, MICs for each antimicrobial agent on the corresponding culture was recalculated and bacteria were subcultured adjusting the compound concentration to 0.5 x MIC. Graph shows the MIC fold change compared to the initially calculated MIC of two independent bacterial cultures, represented by circle and triangle symbols, for each agent. **e)** CCL20-His kills Tet- and Amp-resistant bacterial strains. Parental W3110 bacteria or the three new bacterial strains generated in d, W3110-Amp, W3110-Tet or W3110-CCL20His, were incubated in MHB with Amp, Tet or CCL20-His at a concentration equivalent to the MIC of each compound on parental W3110. Bacteria (5 x 10^+5^ cfu/ml) were incubated at 37°C for 18 h and then bacterial viability in each sample was calculated as in b. Bars represent mean ± SEM % survival data combined from three independent experiments. kDa, kilodalton; MIC, minimum inhibitory concentration.

To study whether *E. coli* develops resistance to CCL20-His, we maintained W3110 bacteria in MHB in the presence of a CCL20-His concentration equivalent to 0.5 x MIC for 2 weeks. For comparison, the conventional antibiotics ampicillin (Amp) and tetracycline (Tet) were also included in this experiment, whose initial MICs, calculated as in Figure 7c, were 5 and 0.5 µg/ml (or 14.3 and 1.1 µM), respectively. On selected days, the MICs of all three compounds were recalculated to assess the evolution of the MIC and to readjust the treatment if necessary to the new MIC. Two separate cultures were initiated and maintained for each antimicrobial compound and analyzed independently. On day 7, the Amp MIC in both cultures treated with Amp increased by > 4-fold and surged to 228 µM (16-fold change) by the end of the experiment (Fig. 7d), indicating that bacteria had become significantly resistant to Amp. Although at a slower rate, the Tet MIC also increased and reached 8.8 and 4.4 µM, an 8- and 4-fold increase, for the two separate Tet-treated cultures, respectively, on day 14. In striking contrast, the CCL20-His MIC remained steady at 20 µM throughout the duration of the experiment (Fig. 7d), indicating that *E. coli* failed to develop resistance against this chemokine. The resulting bacterial cultures after the 14-day conditioning with Amp, Tet, or CCL20-His, were collected, named W3110-Amp, W3110-Tet and W3110-CCL20-His, respectively, and then challenged with a 1 x MIC dose of these 3 compounds. As shown in Figure 7e, consistent with the acquired resistance, Amp and Tet killed the parental W3110 and the W3110-CCL20-His strains but were inactive on W3110-Amp or W3110-Tet, respectively. Of note, the W3110-Tet strain was also resistant to Amp, which is consistent with reports of cross-resistance observed in Tet-resistant bacterial strains^42^. Importantly, CCL20-His killed the parental and all three W3110 conditioned strains (Fig. 7e), which indicates that this chemokine can circumvent Amp and Tet resistance mechanisms to kill bacteria. Furthermore, we found that other antimicrobial chemokines were also able to kill antibiotic-resistant strains effectively. For instance, we found that W3110 and W3110-Amp strains were equally susceptible to CXCL9 (Fig. S7). These results prove that antimicrobial chemokines kill bacteria without triggering microbial resistance and that they inactivate antibiotic resistant microorganisms.

## DISCUSSION

In this study, we demonstrate that chemokine antibacterial activity requires chemokine binding to anionic membrane phospholipids. We show that PG/CL-binding activity is required for microbial killing by chemokines and that all chemokines tested that failed to bind to these membrane anionic phospholipids lacked antimicrobial activity, whereas all those tested that possessed PG/CL-binding activity were antimicrobial. Furthermore, we prove that bacterial membrane CL mediates rapid onset of chemokine bacteriostatic and bactericidal effects. We show that CL-deficient bacteria are more resistant to growth arrest and membrane permeabilization by chemokines, and that antimicrobial chemokines selectively target CL-containing bacteria when these are mixed with CL-deficient bacteria. Importantly, we discovered that bacteria failed to develop resistance against antimicrobial chemokines in our experiments. Since CL is an essential anionic phospholipid in the membranes of most Gram-negative and Gram-positive bacteria, our study provides proof of principle for the development of chemokines as broad-spectrum antimicrobials resistant to antimicrobial resistant mechanisms.

It has been known that bacterial plasma membrane components, and anionic phospholipids in particular, are interaction partners for many classic AMPs^6,8^. However, despite existing evidence of direct bacterial membrane disruption by chemokines^43,44^, this has sometimes been diminished as a secondary or non-specific killing mechanism^21^. Furthermore, the contribution of bacterial membrane anionic phospholipids to this chemokine activity has been overlooked. The first chemokine reported to bind to an anionic phospholipid was CXCL16 with specificity for PS^45^. More recently, in an effort to understand how the highly cationic chemokine MCK-2 encoded by mouse cytomegalovirus mediates cell entry by the virus, we screened an immobilized lipid array and discovered high affinity MCK-2 binding to PS and CL (unpublished data). A subsequent screen with multiple human chemokines revealed that PS and CL binding was common to many but not all chemokines, but that all that bound PS also bound CL^22^. The biological importance of PS to apoptotic cells and apoptotic bodies and the selective localization of CL to bacterial plasma membranes motivated hypotheses for the biological significance of chemokine binding to these phospholipids. In this regard, we have previously reported that chemokine binding to PS may be a ‘find-me’ signal for apoptotic cell clearance by macrophages^22,46^. Our present work provides evidence that binding of chemokines to anionic CL and PG is intrinsic to their mechanism of antimicrobial activity. In addition, our work supports a significant degree of molecular specificity for chemokine-phospholipid binding. For example, we previously reported that chemokines selectively bind CL and PS over other more highly anionic phospholipids, and that some highly cationic chemokines fail to interact with either anionic phospholipid^22^. Thus, chemokine-phospholipid interactions are not solely driven by charge. This agrees with previously published studies that excluded the isoelectric point of chemokines as a reliable predictor of antimicrobial activity^18,19^. Similarly, classic AMPs are generally cationic and amphiphilic but are not limited to these physicochemical properties^4,37^. Here, we demonstrate that the property that may define which chemokines are antimicrobial and which are not is their ability to bind CL-and PG-containing membranes. The precise molecular and chemical properties that allow some chemokines to interact with these anionic phospholipids, and in turn kill bacteria, will require further investigation.

Our findings do not exclude the possibility that, as reported for other AMPs^7^, chemokines may use more than one antimicrobial mechanism, including targeting bacterial proteins or DNA to kill bacteria. In this regard, the transmembrane ATP-binding cassette transporter permease FtsX, the pyruvate dehydrogenase complex (PDHc), and the ABC transport system Opp have been reported to facilitate the killing of *Bacillus anthracis*, *E. coli*, and *Streptococcus pneumoniae*, respectively, by CXCL9, CXCL10 and CXCL11^47–49^. Yet, direct interaction of these chemokines with FtsX, PDHc or Opp has not been demonstrated. A 27-amino acid region in an external loop of FtsX was initially proposed as a putative binding site in *B. anthracis* for CXCL9, CXCL10 and CXCL11 due to its similarity with the N-terminal chemokine-binding domain of CXCR3, the human cellular receptor for these three chemokines^47^.

However, this short region of FtsX shares only 22% amino acid identity with the equivalent region on the N-terminus of CXCR3 and it has been shown that FtsX is dispensable for the killing of *B. subtilis* and *S. pneumoniae* by CXCL10^49^. Moreover, it has been reported that functional FtsX and PDHc rather than their presence are required for CXCL10-mediated killing of *B. anthracis* and *E. coli*, respectively^48,50,51^. FtsX, PDHc and Opp regulate cell division and peptidoglycan synthesis, the conversion of pyruvate to acetyl coenzyme A, and the uptake of oligopeptides involved in nutrition and cell-to-cell communication, respectively, all essential processes for the overall energetic and metabolic state of bacteria^52–56^. Therefore, it may be reasonable to propose that, rather than acting as direct chemokine targets, FtsX, PDHc and Opp may indirectly control bacterial permeability and membrane homeostasis, which ultimately may impair membrane binding and membrane disruption by chemokines or other AMPs. Consistent with this, Opp- and FtsX-deficient bacteria have been shown to be also partially resistant to other membrane active peptides such as nisin and LL-37^49^. Furthermore, it is important to remember that > 20 different chemokines are known to be antimicrobial and that chemokines kill a wide range of bacterial species including *E. coli, Klebsiella pneumoniae, Pseudomonas aeruginosa, S. aureus, S. pyogenes, S. pneumoniae* and others^18,21^. Therefore, a binding site common to all antimicrobial chemokines and conserved across Gram-negative and Gram-positive bacteria, such as CL or PG, may be more likely than the existence of a species-specific bacterial protein ligand for each chemokine. Here, we provide several lines of evidence that support that binding to PG and particularly CL, constitute an intrinsic component of the antimicrobial mechanism of chemokines: i) All antimicrobial chemokines tested in our study (6 out of 6) bind PG and CL whereas all non-antimicrobial chemokines included here (6 out of 6) failed to bind these anionic phospholipids; ii) the binding of CXCL11 to *E. coli* can be blocked by other antimicrobial chemokines, regardless of their different cellular receptors and other biochemical differences, supporting that different antimicrobial chemokines interact with the same bacterial binding sites; iii) bacteria-bound antimicrobial chemokines localize to the poles of the bacterial cell, a region of the *E. coli* plasma membrane where PG and CL phospholipids are known to concentrate^33^; iv) CL-deficient E. coli are more resistant to growth arrest and membrane permeabilization by antimicrobial chemokines; and v) the presence of PG or CL is required for bilayer membrane disruption by antimicrobial chemokines, with CL playing a bigger role than PG in membrane lysis by chemokines but not by other AMPs such as protamine. Although multifunctional molecules like chemokines can be expected to be able to exert different killing mechanisms in different contexts, and it has been reported that some chemokines can act as bifunctional antimicrobial agents^51^, we propose that CL/PG-binding activity of antimicrobial chemokines and derived variants should be tested before establishing membrane-independent bacterial killing mechanisms.

We show here that antimicrobial chemokines can exert bactericidal and bacteriostatic effects and that both antimicrobial effects are promoted by bacterial membrane CL. We found that the absence of CL significantly impaired the ability of antimicrobial chemokines to disrupt phospholipid liposomes and the bacterial plasma membrane and it delayed chemokine-induced bacterial growth arrest. We show that antimicrobial chemokines, but not non-antimicrobial chemokines, are able to lyse liposomes containing a phospholipid composition similar to that of the plasma membrane of *E. coli* (PE/PG/CL) but failed to disrupt liposomes lacking PG and CL. Furthermore, SAXS measurements demonstrated the ability of antimicrobial chemokines to remodel PG-containing liposomal membranes into negative gaussian curvature rich surfaces, a geometry necessary for membrane permeabilization and pore formation^36,37^. On the other hand, understanding the molecular mechanisms by which phospholipid-binding chemokines halt bacterial growth will require further investigation. Anionic phospholipids, and particularly CL, are known to play major roles in bacterial replication by acting as anionic scaffolds in the bacterial plasma membrane for proteins, protein complexes and DNA during cell division^26,57,58^. It is possible that CL-binding chemokines may interfere with the recruitment of these essential elements of the bacterial replication machinery to membrane CL microdomains, ultimately stalling bacterial replication. Another possibility is that similar to the bacteriostatic mechanism of other AMPs such as buforin II or indolicidin^59–61^, antimicrobial chemokines may permeate the bacterial plasma membrane to interact with and destabilize the bacterial genome. Consistent with this hypothesis, some chemokines are known to bind DNA directly^43,62^. In addition, more experimentation will be needed to understand why CL-deficient PG-containing bacteria grow normally early after chemokine challenge but succumb later. In this regard, it would be interesting to investigate the effect of antimicrobial chemokines on PG-deficient bacteria. However, *E. coli* strains lacking PG are not viable unless the major outer membrane lipoprotein Lpp is also deleted^30,63^, which may add uncontrollable effects on fitness and on the mechanical properties of the bacterial membranes^64^, potentially impacting the sensitivity of these strains to membrane active AMPs and making comparisons with PG-containing bacteria problematic.

The results presented here have implications for understanding the role of endogenous AMPs in innate immunity and for the potential development of antimicrobial chemokines as a novel class of antibiotic. To date, of the more than 3,000 discovered AMPs, only nine (daptomycin, colistin, vancomycin, telavancin, teicoplanin, bacitracin, dalbavancin, oritavancin, and gramicidin) have been FDA-approved for clinical use^65,66^. In part, the difficulty in testing an in vivo role for any AMP, including chemokines, relates to the large number of known AMPs and the potential for redundant action^67^. Nevertheless, some existing data support that chemokines may exert antimicrobial effects in vivo. For instance, some chemokines are expressed at high levels in certain barrier tissues without causing inflammation, such as CCL28 or CXCL17 in saliva, and CCL20 in skin and Peyer’s patches^68–71^. CXCL9-depleted mice have been shown to be more susceptible to *Citrobacter rodentium* and *B. anthracis* in a manner that is independent of its receptor CXCR3^44,72^. Although our data show that chemokines need a fairly high concentration to exert bactericidal effects (> 1.2 µM) or reach their MIC (20 µM for CCL20-His), which may be difficult to achieve in vivo, it is possible that endogenous antimicrobial chemokines may act in concert with other AMPs, such as cathelicidins or defensins, or through their bacteriostatic activity, which we show here can be effective at submicromolar concentrations (< 0.3 µM), especially if the target bacteria contain CL. On the other hand, while this concentration-related issue might be easy to overcome in a therapeutic application, a significant challenge for the use of all AMPs in the clinic is their sensitivity to salt concentrations^73^. Although not all tissues and secretions have the same salt content^32^, and a comprehensive analysis of the salt sensitivity of the > 20 different antimicrobial chemokines is lacking, it would be desirable to engineer salt-insensitive chemokine variants. In this regard, certain residue modifications have been known to improve the salt resistance of AMPs^74,75^. Our data support that if these alterations were to be carried out on chemokine sequences, these chemokine variants should preserve their PG- and CL-binding activity. Another limitation of the clinical application of AMPs is their immunogenicity and low stability in vivo^76^. In contrast, recombinant chemokines, as host proteins, can be expected to be non-immunogenic. However, many have a short half-life in blood, potentially driven by serum protease action as well as by scavenging by the erythrocyte chemokine binding protein ACKR1, cognate leukocyte chemokine receptors and glycosaminoglycans^77,78^. Therefore, precise mapping of the CL/PG-binding sites in chemokines may guide generation of antimicrobial chemokine variants with improved bioavailability that could serve as a novel therapeutic approach to treat bacterial infections while preventing the generation of antibiotic-resistant bacteria.

In summary, we have provided evidence that supports specific anionic phospholipid-binding as an important component of the microbial killing mechanism that defines which chemokines are antimicrobial and which are not. We have recently characterized the importance of chemokine interactions with another anionic phospholipid, PS, for phagocyte recruitment in the context of apoptosis^22^. Here, we show that binding to PG and particularly CL is important for microbial killing by chemokines. Together, the present and our previous study demonstrate the potential physiological relevance of these novel chemokine-phospholipid interactions.

## MATERIALS AND METHODS

### Reagents and bacteria strains

Unlabeled recombinant chemokines and hBD3 were purchased from Peprotech (Rocky Hill, NJ) and R&D Systems (Minneapolis, MN), respectively. AZ647-labeled chemokines were acquired from Protein Foundry (Milwaukee, WI).

*S. aureus* strain Wichita (ATCC 29213) was purchased from ATCC (Manassas, VA). *E. coli* strains W3110 and BKT12 were acquired from the Coli Genetic Stock Center at Yale University (New Haven, CT). BKT12 is a CL-deficient mutant of the parental W3110 strain generated by Tan BK *et al*^34^.

### Antimicrobial Assays

*S. aureus* and *E. coli* were grown overnight in tryptic soy broth (TSB) at 37°C. The CL-deficient strain BKT12 was grown in the presence of 50 µg/ml kanamycin. Both, TSB and kanamycin were purchased from KD Medical (Columbia, MD). Stationary cultures were diluted 1:100 in TSB and grown to mid-early log phase (OD^600^ = 0.4 - 0.6). Bacteria were collected by centrifugation (2,500 x g, 5 min) and washed once with antimicrobial assay buffer (AAB, [10 mM Tris-HCl pH 7.4, 1% TSB]). Where indicated AAB was supplemented with 85 mM NaCl (AAB-85). Chemokines and other antimicrobial peptides were incubated with 1 x 10^+5^ cfu of bacteria in 100 µl of AAB for 2 h at 37°C. Bacterial viability was then tested by cfu determination. For this, serial 10-fold dilutions were plated on agar plates in triplicate. Plates were incubated at 37°C overnight and visible cfu were counted manually.

### Liposomes

Phospholipid liposomes were prepared by the extrusion method. All phospholipids used in this study — 1,2-dioleoyl-sn-glycero-3-phosphocholine (PC), 1,2-dioleoyl-sn-glycero-3-phospho-(1’-rac-glycerol) (PG), 1,2-dioleoyl-sn-glycero-3-phosphoethanolamine (PE), 1’,3’-bis[1,2-dioleoyl-sn-glycero-3-phospho]-glycerol (CL), 1,2-dioleoyl-sn-glycero-3-[phospho-rac-(3-lysyl(1-glycerol))] (lysPG) and 1,2-distearoyl-sn-glycero-3-phosphoethanolamine-N-[biotinyl(polyethylene glycol)-2000] (DSPE-PEGbiot) — were purchased from Avanti Polar Lipids (Alabaster, AL). When used for BLI experiments, “PE”, “PG”, “lysPG” and “CL” liposomes contained 30% of the corresponding phospholipid, 65% PC and 5% DSPE-PEGbiot (weight %), and “lpSA” and “lpEC” liposomes consisted of PG/lysPG/CL/DSPE-PEGbiot and PE/PG/CL/DSPE-PEGbiot, respectively, at a 70/20/5/5 ratio (weight %). DSPE-PEGbiot allowed for biotin-mediated immobilization of liposomes onto streptavidin-coated biosensors (SA sensors). When used for competition of chemokine binding or killing of bacteria, “PE”, “PG”, and “CL” liposomes consisted of 70% PC and 30% of the corresponding phospholipids. Phospholipids stored in chloroform were combined at the indicated ratios (for a total of 1 mg) and the solvent was evaporated using a SpeedVac Concentrator (Thermo Fisher Scientific, Atlanta, GA). Dried lipid films were rehydrated for 1 h at room temperature in 0.5 ml of PBS and large unilamellar liposomes were generated by extrusion (>11 passes) through a 0.1 µm pore-sized membrane using the Avanti Polar Lipids mini-extruder. Liposomes were used immediately and prepared fresh for every experiment.

For calcein leakage assays, PE/PG/CL, PE/PG and PE/PC liposomes were generated by extrusion as above by combining the indicated lipids at 70/25/5, 70/30 and 70/30 ratios (weight %), respectively. Dried lipid films were rehydrated for 1 h at room temperature in 0.5 ml of 10 mM Tris-HCl pH 7.4 containing 70 mM calcein (Sigma-Aldrich, St. Louis, MO). Before extrusion, rehydrated phospholipids were subjected to 5 freeze-and-thaw cycles to ensure proper encapsulation of calcein. After extrusion, liposome-encapsulated calcein was purified and separated from any remaining free calcein by size exclusion using Sephadex 50 columns (Sigma) and calcein assay buffer (CAB, [10 mM Tris-HCl pH 7.4, 85 mM NaCl]) as elution buffer. Encapsulated calcein runs fast through the column forming an orange/yellow band whereas free calcein lags as a bright green band. Fractions (0.5 ml) were collected, and all orange fractions were pooled, stored at 4°C, and used within 4 days.

### Biolayer Interferometry

Chemokine binding to liposomes was analyzed by BLI using the Octet RED384 system (Pall ForteBio, Fremont, CA) as previously described ^22^. Before every run, SA biosensors (Pall ForteBio) were hydrated in PBS for 10 min. Then, sensors were equilibrated in PBS for 1 min and liposomes containing DSPE-PEGbiot were immobilized to a final 1-3 nm response. Subsequently, sensors were washed in PBS for 1 min followed by a 5-min incubation in PBS containing 0.05% BSA. Baseline was stabilized in PBS for 5 min, then recombinant chemokine (500 nM in PBS) association was recorded for 500 s, and finally, chemokine dissociation was monitored for 500 s by incubating the sensors with PBS alone. All steps were performed at 1000 rpm and 30°C. In all experiments, background chemokine binding to SA biosensors coated with PC liposomes consisting of PC/ DSPE-PEGbiot (95:5 ratio) was analyzed in parallel and used as reference. Binding to these PC liposomes was subtracted from the binding recorded for every other liposome for each chemokine. Data were analyzed using the Octet Data Analysis software (Pall ForteBio).

### Chemokine binding to bacteria

The binding of AZ647-labeled chemokines was tested by FACS or confocal microscopy. For this, bacteria were grown in TSB to mid-early log phase (OD^600^ = 0.4 - 0.6) and washed once in AAB-85. Then, bacteria (1 x 10^+6^ cfu) were incubated with 0.3 µM of fluorescent chemokine in 100 µl of AAB-85 at 37°C for 20 min. Where indicated, before the addition of the fluorescent chemokine, bacteria were preincubated with 0.3 µM of unlabeled chemokines in 100 µl of AAB-85 at 37°C for 5 min. In addition, to test the effect of different lipids on the chemokine binding to bacteria, in some experiments, AZ647-labeled chemokines were preincubated with 100 µM of PC liposomes containing 30% of PE, PG or CL in 50 µl of AAB-85 at room temperature for 5 min. Then, 1 x 10^+6^ cfu of bacteria in 50 µl of AAB-85 were added to the liposome-chemokine mix and incubated at 37°C for 20 min. All bacterial samples were incubated with AZ647-labeled chemokines in microcentrifuge tubes. Then, samples were washed twice with 400 µl/sample of AAB-85 and bacteria were collected by centrifugation (9,000 x g, 3 min).

For FACS analysis, washed bacteria were co-stained with 10 nM SYTO24 (Thermo Fisher Scientific) in 400 μl of AAB-85 in FACS tubes to distinguish the bacterial cells (SYTO24^+^) from debris (SYTO24^-^). Samples were analyzed in an LSR Fortessa cytometer (BD Biosciences, Chicago, IL) by acquiring 30,000 events at 12 µl/min. Chemokine binding to SYTO24^+^ events was analyzed using FlowJo (BD Biosciences).

For confocal microscopy analysis, washed bacteria were fixed with 2% PFA, immobilized on #1.5 coverslips of 0.17 ± 0.02 mm thickness (Warner Instruments, Hamden, CT) previously coated with 0.1 mg/ml poly-D-lysine (Thermo Fisher Scientific) following the manufacturer’s recommendations, and mounted using Prolong Diamond Antifade mountant with DAPI (Thermo Fisher Scientific) on superfrost plus microscope slides (Fisherbrand, Pittsburgh, PA). Samples were imaged with a confocal laser scanning microscope Zeiss LSM 880 or LSM980 (Carl Zeiss AG, Oberkochen, Germany). We used oil immersion alpha Plan-Apochromat 63X/1.4 Oil Corr M27 objective (Carl Zeiss) and Immersol 518F immersion media (ne=1.518 (30°C), Carl Zeiss). A *z*-stack of images was collected across the entire cell. Identical image acquisition settings, and optimal parameters for x, y, and z resolution were used in all samples from each independent experiment, and representative images for each condition in each experiment are shown with the same display range. Microscopy data processing, analysis, and quantification were done in ImageJ. To quantify bacterial binding, we measured the AZ647 fluorescence intensity of each bacterium at the z-plane containing the highest signal, by generating a region of interest (ROI) around the cell using the oval tool. An equivalent ROI was generated at a region outside the bacterium, considered as background and subtracted from the cell fluorescence intensity. The data were further analyzed and normalized against the control mean using GraphPad Prism 9.

### NAO and chemokine co-staining of *E. coli*

Localization of bacteria-bound CXCL9 was analyzed by Airyscan confocal microscopy in W3110 *E. coli* bacteria co-stained with NAO (Thermo Fisher Scientific). For this, bacterial cultures grown overnight were diluted 1:30 in TSB in the presence of 2 µM NAO and cultured in a lab shaker at 37°C and 220 rpm until OD^600^ ≥ 0.55. Then, bacteria were washed once with AAB-85 and 40 x 10^+6^ cfu were incubated with 0.3 µM CXCL9-AZ647 in 100 µl of AAB-85 at 37°C for 15 min. Bacteria were washed, fixed, immobilized onto poly-D-lysine coated coverslips and mounted on microscope slides, as explained above. A *z*-stack of images was collected across the entire cell on a LSM980 confocal microscope equipped with Airyscan 2 detector (Carl Zeiss) using the super resolution settings in frame mode and optimal parameters for x, y, and z resolution. NAO-524 was imaged with a 488 nm, an MBS 488/561 and SBS SP 550 while a 561 nm argon laser, an MBS 488/561 and a SBS LP 525 was used to image NAO-630. CXCL9-AZ647 was imaged with a 639 nm laser, an MBS 488/561/639 and a SBS LP 640. Airyscan postprocessing was performed using the standard parameters. A line was created along a bacterium and the fluoresce profiles for each channel was generated using ImageJ and further normalized for the minimum and maximum florescence intensity of each independent channel.

### Thin layer chromatography

Phospholipid composition of *E. coli* W3110 and BKT12 was analyzed by TLC as previously described ^34^. Total lipids were extracted from 100 ml of log-phase cultures by the acidic Blight Dyer method. For this, bacterial pellets were resuspended in 2 ml of 0.1 N HCl and mixed with 5 ml of methanol and 2.5 ml of chloroform to generate a single-phase solution. After a 30 min-incubation at room temperature, two phase solutions were created by adding 2.5 ml 0.1 N HCl and 2.5 ml of chloroform. Phases were separated by centrifugation (3,000 x g, 25 min) and the organic lower phase was collected and evaporated using a SpeedVac concentrator (Thermo Fisher Scientific). The dried lipid film was resuspended by sonication in 100 µl of chloroform using a Bioruptor Pico (Diagenode, Denville, NJ). For TLC, samples were spotted on TLC Silica gel 60 plates (Millipore, Bedford, MA) with capillary tubes, and lipids were separated in a TLC developing chamber using a solution of chloroform/methanol/acetic acid (65:25:5, vol/vol) as mobile phase. Lipids were visualized by exposing the plates to iodine vapor by adding a few iodine crystals inside the chamber.

### Time-to-kill assays

The kinetics of chemokine effects on bacterial growth and bacterial killing were analyzed by FACS. For this, 125 µl/well of a bacterial suspension at 10^+6^ cfu/ml in AAB-85 were added in a U-bottom 96-well plate in the presence of buffer alone or the indicated concentrations of chemokine, hBD3, or protamine. Then, plates were incubated at 37°C for 3 h and 25 µl aliquots of each sample were collected at 20, 60, 120 and 180 min for analysis. Aliquots of the inputs before incubation were also collected for determination of the initial number of bacteria (time 0). These 25 µl-aliquots were stained in 400 µl of AAB-85 containing 10 nM SYTO24 and 1.25 µM SYTOX-Orange (both from Thermo Fisher Scientific) in FACS tubes. SYTOX-Orange only permeates dead bacteria with compromised plasma membrane integrity, whereas SYTO24 stains both live and dead bacteria, allowing us to also distinguish bacteria (SYTO24^+^) from debris (SYTO24^-^) during analysis. Tubes were incubated 5 min at RT and events were acquired for 30 s at 12 µl/min in a LSR Fortessa cytometer (BD Biosciences). Data were analyzed using FlowJo (BD Biosciences).

### Calcein leakage assay

The ability of chemokines and other antimicrobial peptides to lyse phospholipid membranes was studied by liposome calcein leakage assays. The fluorescent dye calcein is self-quenched at high concentrations inside liposomes but upon lysis of the liposome and release into the extra-liposomal buffer, calcein regains its fluorescence properties. PE/PG/CL, PE/PG, and PE/PC liposomes encapsulating calcein were prepared and purified in CAB buffer as detailed above (See “Liposomes”). To determine the appropriate volume of liposomes for these assays, calcein release in 10-fold serial dilutions of liposome samples after incubation with CAB containing 0.5% Triton X-100 was first titrated using a FlexStation 3 microplate reader (Molecular Devices, San Jose, CA). Liposome sample volumes causing a 5 to 10-fold increase in the calcein fluorescent signal (ex = 485 nm, em = 515 nm) over baseline were selected for each liposome preparation. Liposomes were first diluted in CAB and 50 µl/well were added in clear bottom black 96-well plates (Greiner, Monroe, NC). Baseline fluorescence was read for 5-10 min in the kinetic mode of the FlexStation 3 reader taking reads every 15-20 s. Then, the read was interrupted, the plate was taken back to the bench and 25 µl/well of buffer alone, chemokine, or protamine prepared in CAB at 3-times the desired final concentration were added. The plate was returned to the microplate reader and the read resumed appending the new read points to the baseline reads. Finally, approximately 30 min later, the read was interrupted again to add 25 µl/well of CAB containing 0.5% Triton X-100 and the fluorescence signal was recorded for additional 5 min to obtain a measurement of the maximum calcein release for each sample. Fluorescence signals were normalized to baseline and the % of calcein release for each sample was calculated as the % fluorescence relative to the maximum fluorescence signal obtained after Triton X-100 addition. The % of released calcein observed for each type of liposome after treatment with buffer alone was subtracted from the corresponding liposome samples treated with the different chemokines or protamine.

### Small angle X-ray scattering experiments with model membranes

Methods used for SAXS experiments and data fitting were based around those that have been previously described^37,79,80^. Liposomes were prepared for SAXS experiments as previously described^37,81^. In brief, lyophilized phospholipids DOPG (1,2-dioleoyl-sn-glycero-3-[phospho-rac-(1-glycerol)]) and DOPE (1,2-dioleoyl-sn-glycero-3-phosphoethanolamine) purchased from Avanti Polar Lipids were dissolved in chloroform stocks at 20 mg/mL. Model membrane lipid compositions were prepared from the lipid stock solutions at a PG:PE 20:80 molar ratio. The lipid composition was evaporated under nitrogen and then desiccated overnight under vacuum to form a dry lipid film, which was resuspended in aqueous 140 mM NaCl, 10 mM HEPES (pH 7.4) to a concentration of 20 mg/mL. Lipid suspensions were incubated overnight at 37°C, sonicated until clear, and then extruded through a 0.2 µm pore size Anopore membrane filter (Whatman) to form small unilamellar vesicles (SUVs). Lyophilized CCL5, CCL19 and CXCL11 powder were solubilized in aqueous 140 mM NaCl, 10 mM HEPES (pH 7.4) and incubated with SUVs at peptide-to-lipid (P/L) charge ratios of 1/6, 1/4, 1/2, 1/1 or 3/2. Samples were hermetically sealed into quartz capillaries (Hilgenberg GmbH, Mark-tubes) for measurements taken at the Stanford Synchrotron Radiation Lightsource (SSRL, beamline 4–2) using monochromatic X-rays with an energy of 9 keV.

The scattered radiation was collected using a DECTRIS PILATUS3 X 1M detector (pixel size, 172 µm), and the resulting 2D SAXS powder patterns were integrated using the Nika 1.50^82^ package for Igor Pro 6.31 and FIT2D^83^. Using Origin Lab software, the integrated scattering intensity I(*q*) was plotted against *q*. Ratios of the measured peak positions were compared with those of permitted reflections for different crystal phases to identify the phase(s) present in each sample. A linear regression through points corresponding to the peaks was used to calculate the lattice parameter, *a*, of each identified cubic phase. For a cubic phase, each peak is represented by a point with coordinates of the assigned reflection (in terms of Miller indices *h*, *k*, *l*) and q. For a cubic phase, 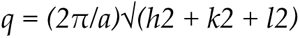. Therefore, the slope of the regression 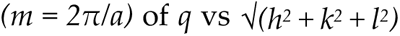 can be used to estimate *a*. The mean negative gaussian curvature (NGC) was estimated as: 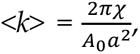 where *A*^0^ and χ are the dimensionless surface area per unit cell and Euler-Poincaré characteristic, respectively, for each cubic phase. *Pm3m*: *A*^0^ = 1.919 and χ= -2, *Im3m*: *A*^0^ = 2.345 and χ= -4, and *Ia3d*: *A*^0^ = 3.091 and χ= -8. For spectra with co-existing *Pm3m* and *Im3m* cubic phases, the ratio of their lattice parameters was noted to satisfy the Bonnet ratio of 1.279.

### Expression and purification of recombinant CCL20-His

Recombinant CCL20-His was expressed and purified using a protocol adapted from previously published methods^84^. pNAN plasmids containing the human *CCL20* coding sequence in frame with a C-terminal short linker (SGGS) and a 6xHis tag, and an ampicillin resistance gene, were purchased from GenScript (Piscataway, NJ). Plasmid (25ng) was transformed into 50 µL of competent Rosetta™ BL21(DE3) pLysS cells via the heat shock method. Cells were streaked onto 150 µg/mL ampicillin agar plates and incubated at 37°C overnight. A single colony was selected for inoculation and grown overnight in 2x Yeast extract Tryptone broth (2xYT) with 150 µg/mL ampicillin. For each liter of 2xYT broth with 150 µg/mL ampicillin, 20 mL of inoculum was added and grown to OD^600^ = 0.6 at 37°C and induced with 0.5 mM IPTG for 6 h before harvesting at 4,000 x *g* and storing overnight at -20°C. For each liter of broth, cells were resuspended in 40 mL of lysis buffer (50 mM Tris HCl pH 8.0, 300 mM NaCl, 0.1 mg/mL benzonase, 10 mM MgCl^2^, and 1 tablet of cOmplete protease inhibitor cocktail [Millipore, Bedford, MA]) and lysed on ice using a Branson 102-C sonifier at 70% power for 3 min (5 s on, 25 s off). After centrifuging at 12,000 *x* g to clarify, the maroon-colored inclusion body pellet was resuspended in 10 mL of Buffer AD (6 M Guanidine HCl, 50 mM Tris, 1 mM tris(2-carboxyethyl) phosphine, pH 8.0) and heated for 30 minutes in a 60°C water bath while passing the suspension through 16- and 20-gauge needles to break up the pellet and dissolve the inclusion bodies. This solution was centrifuged at 24°C for 30 min at 12,000 *x* g to pellet any further insoluble membranes and cellular contents. If centrifugation was insufficient to clarify supernatant, then the solution was filtered at 0.45 µm to achieve an optically clear solution.

For each liter of broth used, 3 mL of Ni Sepharose^TM^ EXCEL resin was used. Resin was washed with 5 column volumes (CV) of milli-Q water, then equilibrated with 5 CV of Buffer AD. The solubilized inclusion body solution was added to resin and allowed to drip by gravity flow, followed by a 10 CV wash with Buffer AD. CCL20-His bound to the column was eluted using 2 CV Buffer BD, (6M Guanidine HCl, 100 mM NaOAc, pH 4.5) twice. Before progressing, the protein was diluted to 1 mg/mL in Buffer BD, quantitated by estimation via its absorbance at 280 nm, and diluted dropwise into 2x volume of cystine/cysteine refolding solution (6.5 mM cysteine, 0.65 mM cystine, 300 mM NaHCO^3^, pH 7.4) and allowed to stir overnight at room temperature. To finalize refolding, the protein solution was dialyzed against 600x volumes of PBS overnight at 4°C, to bring the final guanidine HCl concentration to sub-millimolar concentrations. Refolded CCL20-His was clarified by centrifugation at 12,000 x*g* for 45 min, followed by 0.22 µm filtration. Sample was concentrated using 10 kDa Amicon ULTRA concentrators (Millipore) until an appropriate concentration was reached. Residual guanidine was removed by concentrating and diluting three times with 5x volumes of PBS.

### Minimum inhibitory concentration assay

MICs for chemokines and antibiotics were determined in MHB (Sigma Aldrich). For this, 2-fold serial dilutions (90 µl/well) of CCL20-His, tetracycline and ampicillin were prepared in U-bottom 96-well plates. *E. coli* W3110 strain was grown to mid-early log phase (OD^600^ = 0.4-0.6), and 50,000 cfu of bacteria in 10 µl of MHB were added to each well (final volume = 100 µl/well; final bacteria concentration = 5 x 10^+5^ cfu/ml). Plates were incubated at 37°C for 18 h and MIC was determined as the lower concentration of chemokine/antibiotic showing no evidence of bacterial growth analyzed by luminometry using the BacTiter-Glo Microbial Cell Viability Assay (Promega, Madison, WI).

### Microbial resistance induction assay

To determine the ability of bacteria to develop resistance against antimicrobial chemokines, we monitored over the course of 14 days the change in MIC caused by the exposure of bacteria to a sublethal dose of each antimicrobial agent. For this, *E. coli* W3110 strain was grown in MHB in the presence of CCL20-His, Amp or Tet at a concentration equivalent to 0.5 x MIC. The initial MIC (CCL20, 20 µM; Amp, 14.3 µM; Tet, 1.1 µM) was determined in duplicate by MIC assay. The first day, 50,000 cfu/well of bacteria were cultured in 100 µl of MHB supplemented with the corresponding dose of chemokine/antibiotic in a U-bottom 96-well plate. Two independent bacterial cultures were initiated per treatment and analyzed individually. Bacteria were subcultured every 18 h in 100 µl/well of MHB containing a fresh dose of chemokine/antibiotic. On selected days, MIC was determined for each treatment as explained above, and chemokine/antibiotic dose was adjusted to the new 0.5 x MIC.

### Statistical analysis

Data were analyzed using GraphPad Prism 9. Statistical tests applied for the analysis of each data set are detailed in the corresponding figure legend.

## ACKNOWLEDGEMENTS

S.M.P, S.M, A.Z, K.B, D.G, and P.M.M were supported by the Division of Intramural Research of the National Institute of Allergy and Infectious Diseases (NIAID, NIH). A.B was supported by the Division of Intramural Research of the National Institute of Deafness and Other Communication Disorders (NIDCD, DIR DC000096). J.d.A. was supported by the National Science Foundation (NSF) Graduate Research Fellowship under Grant No. DGE-1650604 and the Ford Foundation. G.C.L.W was supported by NSF DMR2325840. H.A, E.Y.L and G.C.L.W are also supported by the American Heart Association Grant 966662. A.F.D and B.F.V were supported by NIH grant R37AI058072 (B.F.V.).

